# Mechano-osmotic signals control chromatin state and fate transitions in pluripotent stem cells

**DOI:** 10.1101/2024.09.07.611779

**Authors:** Kaitlin P. McCreery, Aki Stubb, Rebecca Stephens, Nadezda A. Fursova, Andrew Cook, Kai Kruse, Anja Michelbach, Leah C. Biggs, Adib Keikhosravi, Sonja Nykänen, Christel Hydén-Granskog, Jizhong Zou, Jan-Wilm Lackmann, Carien M. Niessen, Sanna Vuoristo, Yekaterina A. Miroshnikova, Sara A. Wickström

## Abstract

Acquisition of specific cell shapes and morphologies is a central component of cell fate transitions. Although signaling circuits and gene regulatory networks that regulate pluripotent stem cell differentiation have been intensely studied, how these networks are integrated in space and time with morphological transitions and mechanical deformations to control state transitions remains a fundamental open question. Here, we focus on two distinct models of pluripotency, primed pluripotent stem cells and pre-implantation inner cell mass cells of human embryos to discover that cell fate transitions associate with rapid changes in nuclear shape and volume which collectively alter the nuclear mechanophenotype. Mechanistic studies in human induced pluripotent stem cells further reveal that these phenotypical changes and the associated active fluctuations of the nuclear envelope arise from growth factor signaling-controlled changes in chromatin mechanics and cytoskeletal confinement. These collective mechano-osmotic changes trigger global transcriptional repression and a condensation-prone environment that primes chromatin for a cell fate transition by attenuating repression of differentiation genes. However, while this mechano-osmotic chromatin priming has the potential to accelerate fate transitions and differentiation, sustained biochemical signals are required for robust induction of specific lineages. Our findings uncover a critical mechanochemical feedback mechanism that integrates nuclear mechanics, shape and volume with biochemical signaling and chromatin state to control cell fate transition dynamics.

## Introduction

During early pre- and post-implantation development, the deployment of transcriptional programs is tightly coordinated with morphological transformations of cells to control the maintenance of pluripotency and commitment to the first cell lineages ^1^. When transitioning between states, pluripotent stem cells undergo global changes in their chromatin architecture, resulting in inactivation of pluripotency-specific genes and activation of differentiation-promoting transcriptional programs ^2–4^. These changes in transcriptional programs and their consolidation through epigenetic remodeling are triggered by defined, context-dependent growth factor signals ^5, 6^. For example, in post-implantation primed human embryonic stem cells Fibroblast growth factor (FGF) promotes primed pluripotency whereas transforming growth factor-β (TGF-β) signaling regulates both naïve and primed pluripotency^7, 8^. In naïve pluripotent cells FGF regulates hypoblast commitment^9–11^. Besides these key biochemical inputs, mechanochemical signals imparting from the extracellular matrix and the actomyosin cytoskeleton have been shown to be important for proper cell fate cell selection and differentiation of various pluripotent stem cells ^12–14^

During the transition from pluripotency to the commitment to specific cell lineages, embryonic stem cells also undergo distinct morphological and mechanical changes as the cells transit into their specialized shapes and structural configurations ^13, 15, 16^. Studies in other cellular systems have shown that morphological and volumetric changes also have the potential to regulate gene expression and metabolic activity through mechanosensitive signaling pathways such as the YAP pathway^17–19^. Besides activating membrane-mediated cascades of intracellular signaling, extrinsic mechanical forces and osmotic pressures also have the ability to directly deform the cell nucleus, driving the adaptation of nuclear morphology phenotypes and genome architecture ^20–24^.

Beyond these effects of mechanical forces on transcription, the role of dynamic mechanical properties of the nucleus and chromatin itself in gene regulation is less understood. On the scale of the entire nucleus, chromatin behaves like an elastic solid, but on the scale of single chromatin domains it displays liquid-like properties ^25–28^. As these rheological processes influence the dynamics of molecular processes, dynamic changes in chromatin mechanics for example through changes in chromatin fiber interaction by multivalent chromatin-binding proteins such as chromatin remodelers, have the potential to regulate genome activity ^29, 30^. Moreover, on a larger scale, mechanisms like remodeler- and ATP-dependent chromatin stirring may facilitate protein access to dense chromatin regions, though the physiological relevance on such mechanisms remains unclear ^31^. Thus, how cell shape transitions, nuclear mechanical and volumetric properties, and their influence on chromatin and gene expression are coupled during lineage transitions presents a fundamental open question.

Here, we use human induced pluripotent stem cells (hiPSCs) to investigate how morphological and osmotic changes are integrated with biochemical signaling and gene regulatory networks to gate cell fate transitions. Further, we used donated human blastocyst stage embryos to address if similar mechanisms are applied in embryonic fate decisions more broadly. We find that cell fate transitions in both models are associated with a decrease in nuclear volume. Mechanistic studies in the primed hiPSCs reveal that removal of growth factors that maintain primed pluripotency generates mechanical and osmotic stresses that act in concert to modulate nucleoplasmic viscosity and nuclear stiffness. Consequently, macromolecular crowding and biomolecular condensation prime chromatin for a subsequent cell fate transition. On short timescales, permissive biochemical conditions allow mechano-osmotic forces to enhance the efficiency of fate conversion by inducing chromatin-level de-repression. We thus propose that mechano-osmotic reprogramming of the nuclear environment tunes differentiation efficiency by lowering the energy barrier for cell fate transitions.

## Results

### Exit from primed pluripotency is associated with mechano-osmotic remodeling of the nucleus

To address the relationship between cell morphology and cell fate specification we used hiPSCs (from the Allen Institute) that represent a post-implantation, primed pluripotent stem cell state as a model ^32^ ^33^. Morphometric analyses revealed that differentiation of these hiPSCs from primed pluripotency into the three germ layers was accompanied by a reduction in nuclear volume and nuclear flattening (Fig. 1a-c; Supplementary Fig. S1a-d), paralleling observations of nuclear flattening in mouse embryonic stem cells exiting naïve pluripotency ^34^.

**Figure 1:**
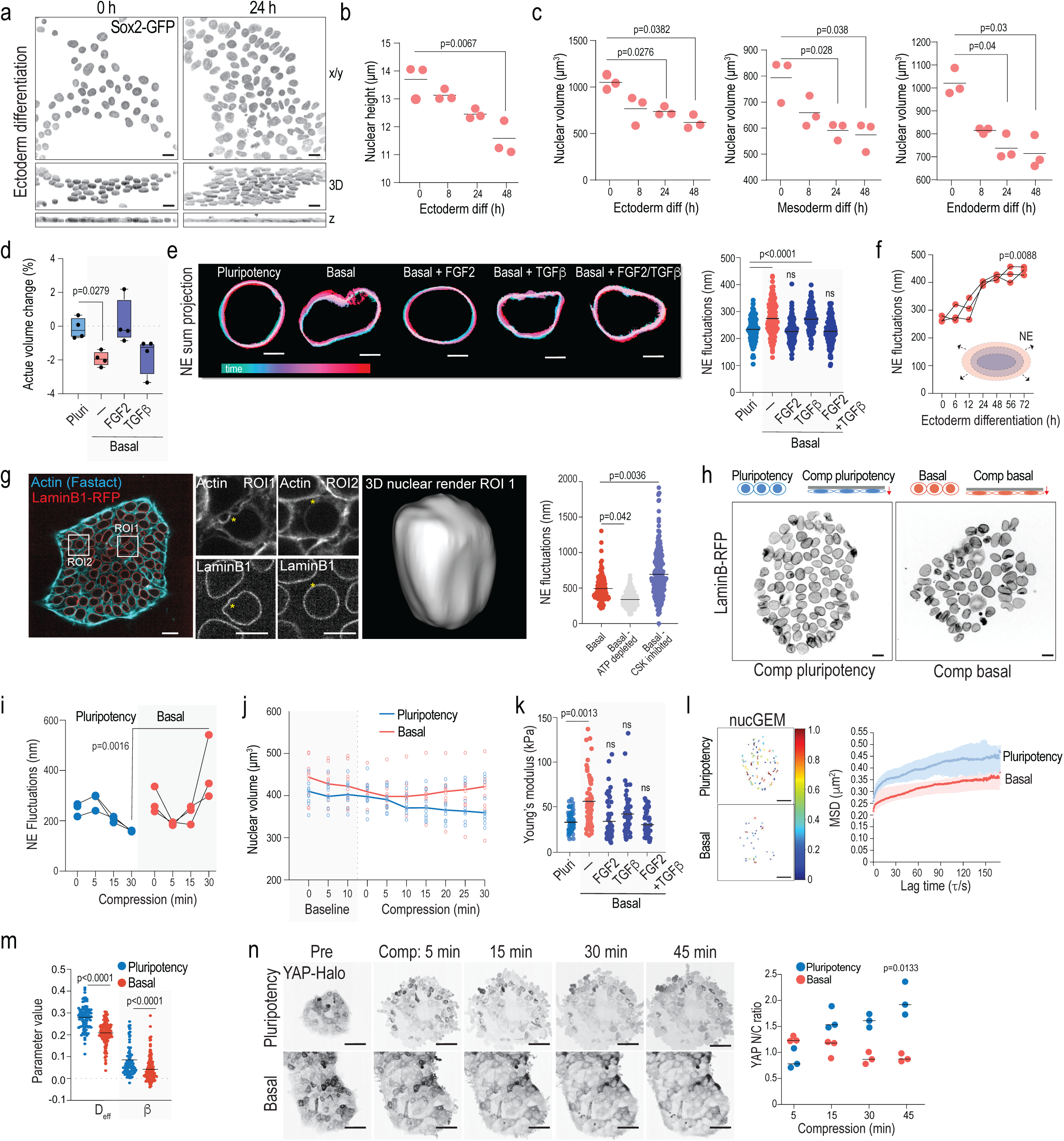
Exit from primed pluripotency is associated with mechano-osmotic remodeling of the nucleus. **(a)** Representative top views (x-y), 3D reconstructions and cross sections (z), of Sox2-GFP-tagged hiPSCs undergoing ectodermal differentiation for the indicated time points (scale bars 15 µm). **(b)** Quantification of nuclear height from hiPSCs undergoing ectodermal differentiation for the indicated time points (n= 3 independent experiments with >600 nuclei/condition/experiment; ANOVA/Dunnett’s). **(c)** Quantification of nuclear volume from hiPSCs undergoing tri-lineage differentiation for the indicated time points (n= 3 independent experiments with >600 nuclei/condition/experiment; ANOVA/Dunnett’s). **(d)** Quantification of change in nuclear volume upon exposure to culture medium/growth factors indicated (n=4 independent experiments with >60 nuclei/condition/experiment; Kruskal-Wallis/Dunn’s). **(e)** Representative projections of nuclear envelope movements as a function of time upon pluripotency factor removal and adding back specific growth factors. Note that removal of pluripotency factors triggers immediate fluctuations that can be rescued by adding back TGF-β and FGF2 (scale bars 5µm; n= 3 independent experiments with >140 nuclei/experiment; ANOVA/Kruskal-Wallis). **(f)** Quantification of rapid nuclear envelope fluctuations in iPSCs undergoing ectodermal differentiation for time points indicated (n= 3 independent experiments with >120 nuclei/condition/experiment; Kruskal-Wallis/Dunn’s). **(g)** Representative snapshots of live imaging movies of Lamin-B-RFP tagged iPSC in basal medium, stained with FastAct to label Actin. Note perinuclear actin rings surrounding nuclei and actin-rich bleb like structures with corresponding nuclear deformation (scale bars 20 and 10 µm; images representative of 5 movies). **(h)** Quantification of nuclear fluctuations from cells in basal medium with or without ATP depletion. Note attenuation of fluctuations upon ATP-depletion (n>150 nuclei/condition pooled across 3 independent experiments; ANOVA/Kruskal Wallis). **(i, j)** Schematic of experimental outline, representative images (i), and quantification (j) of nuclear envelope fluctuations in cells compressed in pluripotency or basal medium for time points indicated. Note decreased fluctuations pluripotency condition and an increase in basal medium (scale bars 10 µm; n> 150 nuclei/condition pooled across 3 independent experiments; ANOVA/ Kruskal-Wallis). **(k)** Quantification of nuclear volume evolution over time from of Sox-GFP-tagged hiPSCs live imaged directly after a media change into pluripotency or basal medium, followed by compression. Line represents median volume and individual dots are average colony volumes at indicated timepoints. (n= 10 colonies/condition pooled across 6independent experiments). **(l)** AFM force indentation experiments of iPS cell nuclei within 20 min of media switch. Note increased elastic modulus of cells in basal media conditions, restored by adding FGF2 (n > 70 nuclei/condition pooled across 5 independent experiments; ANOVA/Kruskal-Wallis). **(m)** Representative tracks of nucGEM particles. Colors represent average rate of diffusion per tracked particle (scale bars 5 µm). **(n, o)** Quantification of mean squared displacement versus lag time per nucGEM particle (h) and nucGEM diffusion and diffusivity exponent β (n=3 independent experiments with >140 nuclei/experiment; ANOVA/ Kruskal-Wallis). **(p)** Representative snapshots of live imaging and quantification of HALO-tagged endogenous YAP localization in cells compressed to 5 µm height in pluripotency or basal medium. Note YAP nuclear entry in pluripotency condition but not in basal medium upon compression (scale bars 30 µm; n= 3 independent experiments with >100 nuclei/experiment; ANOVA/Friedman).

To determine if these morphological transitions occur before or after loss of pluripotency, we analyzed the immediate response of nuclear volume to removal of FGF2 and TGF-β1 factors which maintain primed pluripotency in this system ^35^. Strikingly, replacing the pluripotency-maintenance medium with basal medium lacking these growth factors resulted in rapid reduction of nuclear volume within 15 min of media change, despite unchanged media osmolarities (Supplementary Fig. S1e). This effect was mainly driven by the absence of FGF2 signaling as adding FGF2 to the medium prevented nuclear volume decrease, while adding TGF-β1 had a less prominent impact on volume restoration (Fig. 1d). To investigate the mechanisms of this rapid, minute-scale nuclear volume change, we first analyzed nuclear envelope mechanics by quantifying rapid nuclear fluctuations using fast imaging of endogenously tagged LaminB-RFP iPSCs ^36, 37^. Nuclear envelope fluctuations increased immediately following the removal of pluripotency-maintaining growth factors, appearing within minutes of the medium exchange (Fig. 1e). Importantly, similarly to the decrease in nuclear volume, nuclear fluctuations were specifically controlled by growth factors (Fig. 1d). While adding back TGF-β1 further enhanced rapid local high amplitude fluctuations, FGF2 restored the taut nuclear envelope phenotype observed in pluripotency conditions. Adding both factors together fully restored nuclear dynamics to the pluripotent cell state, suggesting that growth factor signaling drives nuclear mechano-remodeling (Fig. 1d). To validate that nuclear fluctuations are associated with cell state transitions, we quantified nuclear fluctuations during biochemical differentiation of hiPSC colonies towards the three germ layers and noted a gradual increase of nuclear fluctuations as hiPSCs lost markers of pluripotency and began expression markers of germ layers (Fig. 1f; Supplementary Fig. 1d).

We next set out to uncover the driving forces behind observed fluctuations and test if they are connected to functional morphological changes during fate transitions. Given the rapid timescale and vast experimental evidence between growth factor signaling and cytoskeletal remodeling ^38^, we hypothesized that nuclear envelope fluctuations may result from the perinuclear cytoskeleton confining and actively deforming the nucleus. Indeed, live imaging of actin dynamics revealed presence of a taught, perinuclear actin ring encapsulating the nuclear envelope (Fig. 1g). Intriguingly, we further noted the presence of dynamic actin structures that originated from the extracellular space and actively deformed the nucleus on the same spatiotemporal scales observed for nuclear fluctuations (Fig. 1g).

To understand if the observed cytoskeletal dynamics trigger nuclear envelope fluctuations, we disrupted the cytoskeleton using a combination of cytochalasin D (actin) and nocodazole (microtubules) and quantified nuclear fluctuations. These analyses revealed amplification of fluctuations with pharmacological inhibitors of actin and microtubules suggesting that the perinuclear cytoskeleton restricts and confines the nucleus (Fig. 1h). We then set out to test whether pushing forces emanating from chromatin rearrangements might counteract these cytoskeletal constraints and cause fluctuations, as previously suggested ^36^ . To test this, we depleted cellular ATP to block energy-dependent cellular activities including dynamic chromatin remodeling and found that the lack of ATP significantly reduced nuclear envelope dynamics, supporting an active remodeling process (Fig. 1h).

Our data thus far suggested that removal of primed pluripotency-maintaining growth factors maintaining induce rapid nuclear flattening and volume reduction, and increase nuclear envelope fluctuations in hiPSCs. To investigate the link between flattening and nuclear envelope mechanics, we mimicked the cytosleketal confinement using a compression bioreactor to flatten nuclei by confining hiPSC colonies to a height of 5 µm which amounted to approximately 30-40% decrease in nuclear height (Fig. 1i). We found that nuclear flattening affected nuclear fluctuations, but the effect was determined by the biochemical signaling environment. Specifically, in pluripotency-promoting media, compression reduced nuclear envelope fluctuations and increased nuclear envelope tautness. In contrast, in basal media lacking pluripotency growth factors, compression increased nuclear wrinkling and amplified nuclear envelope fluctuations (Fig. 1i, j), consistent with the role of growth factors determining nuclear mechanics. Intriguingly, growth factors also controlled the rates of nuclear volume loss triggered by extrinsic compression. In the absence of pluripotency growth factors hiPSC exhibited rapid volume loss which was followed by a rapid recovery, while in pluripotency medium, the volume decreased gradually, peaking only after 30 min of confinement (Fig. 1k).

The context-dependent effect of extrinsic compression on nuclear fluctuations and volume change suggested that pluripotency factors determine the nuclear mechanophenotype. To corroborate this, we measured nuclear/chromatin composite stiffness by atomic force microscopy (AFM)-mediated force indentation spectroscopy^27^. Removing pluripotency factors triggered nuclear stiffening within 5 minutes of the perturbation (Fig. 1l). Adding FGF2 to the basal medium prevented this stiffening, while TGF-β1 had milder effect. Both factors together restored the pluripotent nuclear mechanical state (Fig. 1l). Similar nuclear stiffening was also observed when nuclear volume loss was induced by transient hypertonic shock (Supplementary Fig. S2a, b). We thus asked if this rapid stiffening was due to changes in nucleoplasmic/chromatin viscosity caused by volume loss. To measure the rheological properties of the nucleoplasm, we utilized nuclear Genetically Encoded Multimeric nanoparticles (nucGEMs)^39^. Analysis of the mean square displacement of these particles showed reduced diffusion and increased confinement in basal conditions compared to pluripotency medium (Fig. 1n-o). Taken together, these findings indicate that withdrawal of pluripotency-maintenance growth factors leads to immediate changes in the nuclear mechanophenotype: increase in nuclear stiffening and induction of osmotic stress, which together increase macromolecular crowding of the nucleoplasm.

Recognizing the role of mechanical forces in embryonic cell fate specification ^13, 24, 34, 40, 41^, we hypothesized that nucleomechanical changes might affect hiPSC mechanosensitivity. To test this, we turned to the activation of the yes-associated protein (YAP), as this transcriptional regulator is known to integrate inputs from cell mechanics and metabolism to control cell fate segregation, pluripotency and germ layer specification in a dynamic, dose-dependent manner ^19, 42–46^. First, we monitored nuclear YAP levels during hiPSCs differentiation into the three germ layers and observed a progressive loss of active YAP within 8 hours of differentiation, analogous to the timeline of nuclear volume changes (Supplementary Fig. S2c). We thus hypothesized that the growth factor-controlled NE mechanics could be gating YAP activity. This notion was supported by the anti-correlated kinetics of YAP and NE fluctuations upon biochemically triggered exit from primed pluripotency (Supplementary Fig. S2d). To determine if YAP nuclear entry is regulated by nuclear tautness/flattening or by volume loss, we compared YAP dynamics in response to a hyperosmotic shock and compression. Live imaging of iPSCs with endogenously Halo-tagged YAP revealed a salt-and-pepper like pattern of active YAP in the nucleus at steady state. Hypertonic stress did not substantially alter this localization pattern (Supplementary Fig. S2e), indicating that YAP is not controlled by nuclear volume loss. In contrast, compression in the presence of pluripotency growth factors increased YAP nuclear entry, yet the same mechanical manipulation in the absence of these growth factors failed to strongly activate YAP, indicating that growth factor-controlled nuclear fluctuations gate YAP activation (Fig. 1q). Consistently with such thresholding activity, subjecting iPSCs to a greater deformation (3 μm instead of 5 μm) resulted in YAP activation also in the absence of growth factors (Supplementary Fig. S2f).

Finally, we asked if the observed mechano-osmotic remodeling of the nucleus was restricted to primed iPSCs or relevant to embryonic cell fate transitions more broadly. To this end we turned to the emergence of the first cell types in naïve pluripotency ^32^. The first embryonic cell lineage segregation to either trophoblast or inner cell mass (ICM) is observed when the embryo proceeds from the morula to early blastocyst stage ^47–49^. The ICM then segregates into epiblast and hypoblast where the epiblast expresses pluripotency-associated transcription factors OCT4 and NANOG, whereas the hypoblast comprises cells that express SOX17, GATA6, GATA4 and PDGFRA ^32^. To obtain insights into the morphological transitions that accompany and control early human development, we investigated the shapes of the ICM cell nuclei during their first lineage transitions. For this, we obtained surplus human embryos (5 embryos) donated for research after artificial reproduction cycles. These embryos were carefully morphologically staged, after which whole mount immunostainings were performed of the epiblast marker NANOG and the hypoblast marker GATA6, together with DAPI to mark chromatin and Lamin-B1 to mark the nuclear lamina (Supplementary Fig. 2g). Using 3D segmentation, we observed that the nuclei of GATA6-high cells had reduced volumes and increased surface to volume ratios compared to NANOG-high cells within the inner cell mass, indicative of flattening or deformation (Supplementary Fig. 2h). As in the primed iPSCs, actin structures were visible in the blastocyst stage human embryos, where they closely correlated with nuclear deformation (Supplementary Fig. 2i), indicating that mechano-osmotic forces associate with lineage transitions in the early human embryo.

Collectively, these experiments demonstrated that cell fate transitions in pluripotent cells are associated with increased active nuclear envelope fluctuations driven by a composite effect of actin dynamics-mediated confinement and intranuclear forces, increased nucleoplasm crowding/viscosity, increased nuclear stiffness, and dampened nuclear mechanosensitivity. The results further showed that these mechano-osmotic changes are directly controlled by the coordinated action of pluripotency-maintaining growth factors.

### Nuclear flattening primes chromatin for spontaneous differentiation

We next asked whether the observed mechano-osmotic changes of the nucleus affect cell fate transitions. To test this, we flattened cells in pluripotency medium for 5 or 30 min using the compression bioreactor (Fig. 2a) and characterized immediate cellular responses and their potential heterogeneity by measuring genome-wide chromatin accessibility and transcription (single cell multiome ATAC and RNA sequencing; 10xGenomics platform). We also examined the long-term reversibility/sustainability of potential changes by analyzing a recovery condition 24 h after the 30 min compression. Already after 5 min of compression, we detected changes in chromatin accessibility and gene expression that became more prominent after 30 min of compression (Fig. 2b; Supplementary Fig. S3a). This response was robust and stereotypic, as evidenced by a cluster consisting almost exclusively of 30 min compressed cells predominantly in the ATACseq dataset (Fig. 2b), whereas 24 h recovery condition clustered with uncompressed cells.

**Figure 2:**
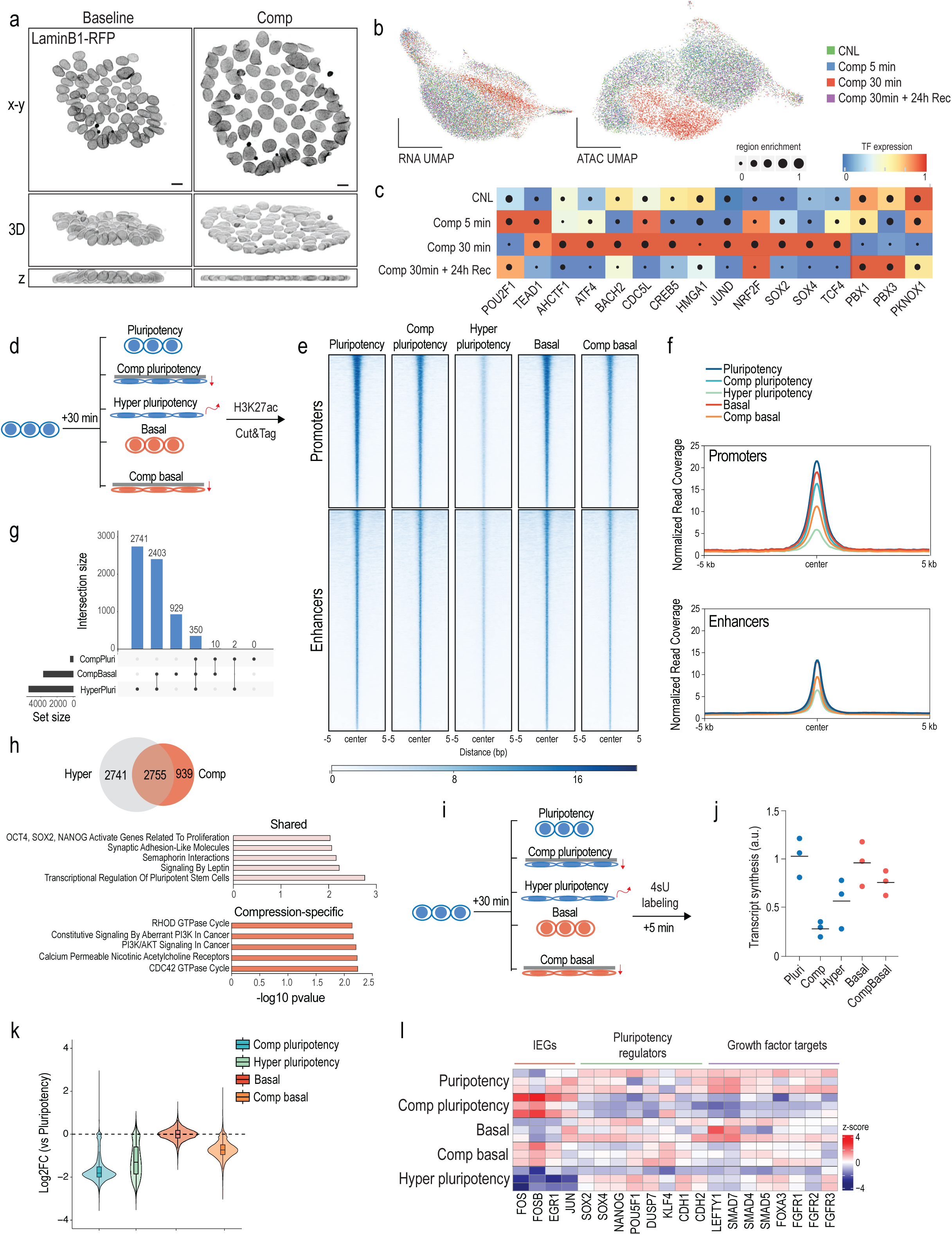
Nuclear flattening primes chromatin for spontaneous differentiation. **(a)** Representative top views, side views and 3D reconstructions of LaminB1-RFP-tagged hiPSCs subjected to compression (Scale bars 10 µm; images representative of 6 independent experiments). **(b)** Uniform manifold approximation and projection (UMAP) of scRNA- and scATAC-seq from hiPSCs subjected to compression for timepoints indicated **(c)** Heatmap of predicted regulons enriched in compressed cells from SCENIC+ analyses of the multiome data. **(d)** Schematic of experimental outline for genome-wide mapping of H3K27ac changes. **(e, f)** Heat map (e) and metaplot (f) analysis of mean H3K27ac levels at active promoters and predicted active enhancer regions. Note reduction in H3K27ac enrichment at promoters across all conditions and at enhancers in cells compressed in basal medium or exposed to hypertonic shock. **(g)** UpSet plot showing an overlap of enhancers decommissioned in compression and hypertonic shock conditions. **(h)** Venn diagram and Reactome pathway enrichment of compression-specific and shared decommissioned enhancers as defined in (g). **(i)** Schematic of experimental outline for quantification of the nascent transcriptome. **(j)** Quantification of RNA synthesis across conditions. Note reduced synthesis across all conditions compared to pluripotency medium condition. **(k)** Quantification of total changes in nascent RNA production across conditions relative to the pluripotency medium condition. **(l)** Heatmap of altered nascent RNA levels of relevant transcripts from TTseq. Note increased levels of immediate early genes specifically in cells compressed in pluripotency medium while key pluripotency and growth factor regulators are repressed.

Overall chromatin accessibility appeared reduced with 5131 regions losing accessibility and only 157 regions gaining accessibility at 30 min. To elucidate the regulatory circuits between the observed chromatin changes, transcription factors and gene expression, we applied SCENIC+ analyses^50^ to detect integrated gene regulatory networks after 5 or 30 min of compression and after 24 h recovery following 30 min compression. The co-regulator of the mechanosensitive transcription factor YAP, TEAD1, and the mechanosensitive POU2F1 (Oct1)^19^ were detected as upregulated regulons already at 5 min of compression (Fig. 2c), while the regulatory networks downstream of key master pluripotency transcription factors including SOX2, SOX4 and TCF4, as well as key master regulators of stress responses JUND, ATF4, and NRF2 peaked a bit slower after 30 min of compression (Fig. 2c). Overall, chromatin accessibility and gene expression changes appeared largely reversible as the 24 h recovery condition closely resembled the control condition (Fig. 2b, c). Notably, no induction of apoptosis/necrosis genes or effects on cell cycle were observed (Supplementary Fig. S3b, c), indicating that compression did not trigger damage/cell death or influence cell cycling.

To further investigate the relationship between nuclear mechanics and pluripotency, we used chromVAR ^51^ to predict transcription factor ‘activity’ based on the enrichment of binding motifs for increasingly accessible chromatin regions following compression-driven nuclear flattening. The strongest signature came from TEAD1-4 targeted motifs, as well as the SOX, POUF, and SIX families of TFs involved in gastrulation, indicative of chromatin priming towards exit from pluripotency (Supplementary Fig. S3d). Differential gene expression analyses further confirmed that genes involved in the regulation of the actomyosin cytoskeleton including known YAP target genes (AMOTL1, TAGLN, CCN2), as well as genes involved in heat stress and osmotic stress (HSPA1A, HSPA1B, FOS) were upregulated at 30 min (Supplementary Fig. S3e). However, in contrast to chromatin profiling, differential gene expression analysis further revealed that genes involved in pluripotency such as SOX2, OCT4, and DUSP7 were largely unchanged or even slightly upregulated upon 30 min of compression (Supplementary Fig. S3f, f). Thus, whereas the chromatin profile indicated priming of chromatin towards exit from pluripotency, activity of predicted transcription factors such as YAP/TEAD and RNA profiling described a more complex transcriptional state that included a signature of reinforcement of pluripotency-associated transcriptional circuitry.

As nuclear flattening by mechanical compression triggered chromatin priming towards lineage transition but no sustained fate transition, we reasoned that the pluripotency-promoting growth factors were preventing exit from pluripotency. To test this, we quantified genome-wide changes in promoter and enhancer states by profiling the epigenetic mark of active enhancers, H3K27ac, in pluripotency or basal media conditions lacking FGF2 and TGF-β1 with or without compression for 30 min (Fig. 2d; Supplementary Fig. S4a). To separately evaluate the impact of volume loss and the associated osmotic stress response on enhancer landscape, we included a hypertonic shock condition. Intriguingly, removal of pluripotency factors (basal medium) or compression in pluripotency medium triggered a decrease in H3K27ac at promoters but had no strong effect on enhancers (Fig. 2e, f; Supplementary Fig. S4a). In contrast, compression in basal medium caused a strong reduction of H3K27ac both at promoters and enhancers, similar to hypertonic shock (Fig. 2e, f). The decommissioned enhancers in these conditions showed high overlap, with many of the enhancers affected by compression in pluripotency conditions also changed in basal media compression and hypertonic shock conditions (Fig. 2g, h). Annotating the enhancers by assigning them to the nearest gene revealed that the shared enhancer changes were enriched for key pluripotency genes such as SOX2, LIN28A, and ZIC3 (Fig. 2h). In contrast, the compression-specific decommissioned enhancers were enriched for cytoskeletal regulators and PI3Kinase signaling, whereas hyperosmotic-specific enhancers showed enrichment for FOXO targets (Fig. 2h. Supplementary Fig. S4b).

The strong effects of compression in de-commissioning promoters and enhancers, similar to hypertonic stress, suggested a global impact on transcription. To quantify this and to understand the immediate transcriptional responses to mechanical deformation in different contexts, we analyzed the nascent transcriptome (transient transcriptome sequencing; TTseq^52^) after 30 min exposure to pluripotency-promoting culture conditions or basal medium, with or without compression, and in response to hypertonic shock as control for osmotic regulation (Fig. 2i). Quantification of RNA synthesis from nascent RNA labeled during the last 5 min of 30 min of exposure indeed revealed reduced levels of global transcription upon compression and hypertonic shock (Fig. 2j, k).

Surprisingly, this transcriptional repression was strongest in compression in pluripotency medium. Further analysis of differential gene expression between compressed conditions revealed decreased expression levels of more than 15,000 genes in the compressed pluripotency condition compared to pluripotency medium control and over 8,000 genes when compared to compression in basal medium, including mediators of differentiation (Supplementary Fig. S5a). Meanwhile, although transcription was overall strongly repressed in compressed cells in pluripotency medium, classical differential gene expression analysis of the nascent transcriptome revealed strong activation of immediate early genes (IEGs; eg JUN, FOS, EGR1), a class of genes that are rapidly and very transiently induced in response to stress or growth factor stimuli in a strictly time and dose-dependent manner ^53–55^ (Fig. 2l; Supplementary Fig. S5b). This was consistent with the multiome analyses showing increased expression of JUN/FOS targets in this condition (Supplementary Fig. S3e). Such strong IEG induction was not apparent upon 30 min compression in basal medium (Fig. 2d-f) again pointing to biochemical context-dependency of nuclear mechanotransduction in iPSCs.

The notion of pluripotency-promoting growth factors gating the transcriptional response to mechanical flattening was supported by the kinetics of the stress-sensitive kinase p38 MAPK. While p38 was strongly activated by compression in basal medium already at 5 mins and downregulated at 30 mins, coinciding with transcriptional repression, p38 activation and transcriptional repression only became apparent in pluripotency medium at 30 min of compression, the timepoint of nascent RNA profiling (Supplementary Fig. S5c, d). The slower dynamics of p38 activation was consistent with the kinetics of nuclear volume loss that peaked at 30 min in this condition (Fig. 1l). Importantly, adding FGF2 into the basal medium suppressed p38 activation by compression, whereas TGF-β1 did not have such a strong effect, further confirming that FGF2 signaling controls the mechano-osmotic properties of the nucleus and its response to deformation (Supplementary Fig. S5d).

Collectively these data indicated that mechanical deformation of the nucleus has two distinct components: a mechanical component that impacts YAP activity and cytoskeletal/extracellular matrix gene expression, and an osmotic stress component that is sufficient to induce a chromatin state priming towards exit from primed pluripotency in hiPSCs even in the presence of pluripotency-promoting growth factors. However, in pluripotency medium this chromatin priming effect is weaker and transient, and while promoters are strongly impacted by compression, enhancers remain largely unchanged thus locking existing cellular fates. Further, pluripotency growth factor-mediated signals determine the mechano-osmotic state of the nucleus and the associated dynamics of the transcriptional repression in response to nuclear flattening, and removal of these pluripotency factors enhance the osmotic component of deformation to efficiently remodel enhancers.

### Mechano-osmotic signals control kinetics of lineage commitment

To directly address the role of mechano- and osmosensitive enhancer remodeling on long-term transcriptional memory and lineage commitment, we performed bulk-RNAseq 24 h post recovery from 30 min compression, either in pluripotency maintenance conditions or in basal medium, comparing these scenarios to hypertonic shock (Fig. 3a). Principal component analysis revealed that either transient hypertonic shock or compression each triggered distinct long-term transcriptional responses under conditions of pluripotency. In contrast, in basal media conditions which alone promoted a transcriptional signature of spontaneous differentiation, compression and hypertonic shock led to a highly correlated transcriptional state lacking a specific signature of mechanotransduction in the compressed condition (Fig. 3b). This further supported the notion that growth factor signaling directly gates the balance between the mechanical and osmotic components of compression. Further, while in the pluripotency condition the long-term transcriptional response to transient compression showed a gene expression signature of FOS and STAT3 transcriptional targets and enrichment for cytoskeletal genes, consistent with the dominance of the mechanoresponse and YAP activity in this condition (Fig. 3c-e), in the basal media conditions transient compression led to further upregulation of a gene regulatory network of “non-lineage-specific” differentiation and downregulation of pluripotency genes compared to basal media alone (Fig. 3c-e; Supplementary Fig. S6a). A similar activation of differentiation gene signatures was also observed in the hypertonic condition in the basal medium condition (Supplementary Fig. S6a). The osmosensitive metallothionine gene family ^56^ was most upregulated in cells transiently compressed in basal media conditions (Fig. 3c), further indicating that the long-term transcriptional effects of this condition closely resemble hypertonic shock. Collectively, the gene activation profiles indicated that withdrawal of pluripotency factors, together with nuclear volume loss, led to acceleration of differentiation.

**Figure 3:**
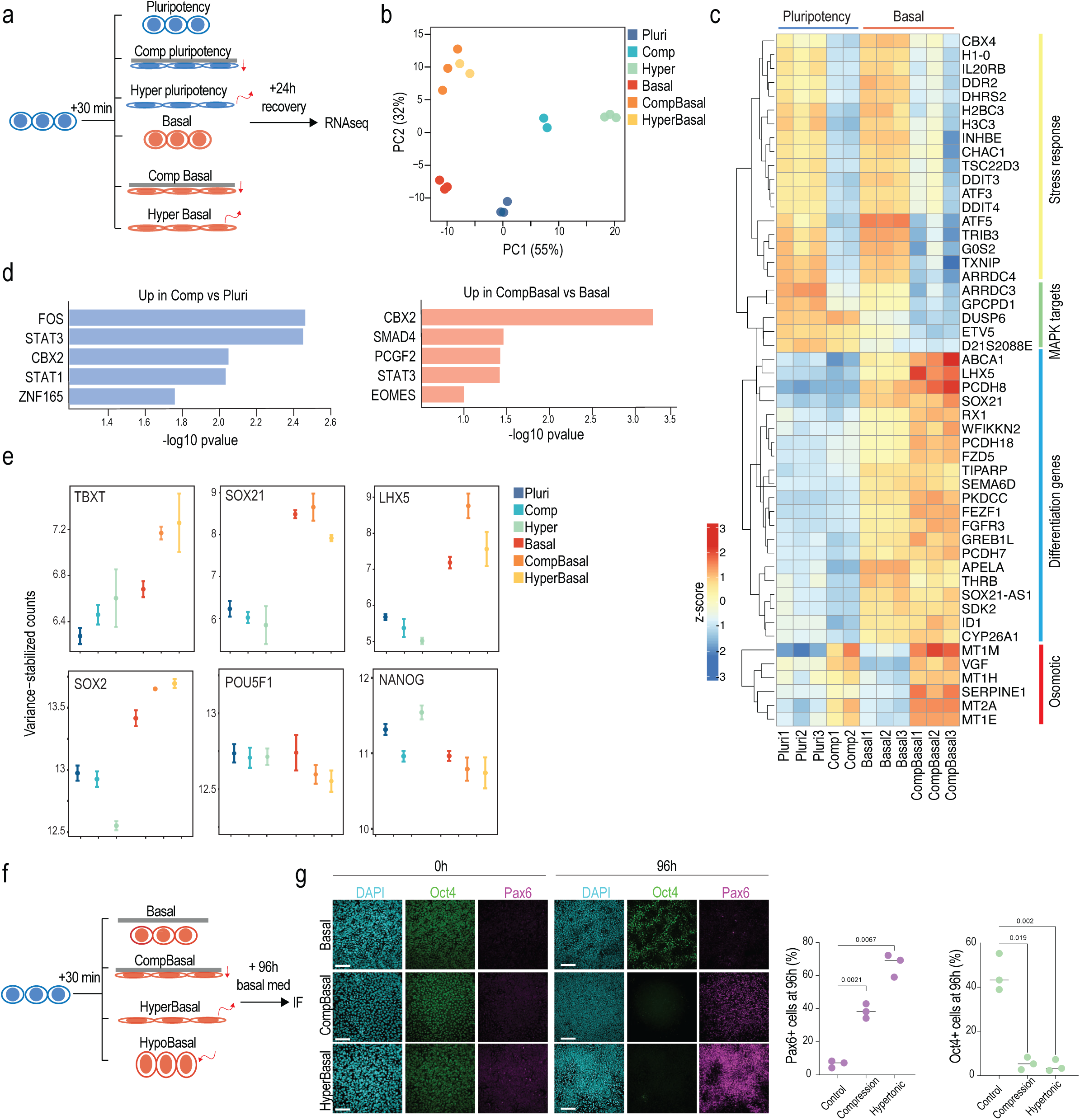
Mechano-osmotic signals control kinetics of lineage commitment. **(a, b)** Schematic of experimental outline (a) and PCA plot (b) of bulk RNA sequencing in cells subjected to compression or hypertonic shock and recovery in the indicated media conditions. Note divergence of transcriptome of hypertonic shock and compression in pluripotency medium and convergence in basal medium. **(c)** Heatmap of top variable genes from bulk RNA seq in conditions indicated. Note increase in differentiation gene expression in cells compressed in basal medium. **(d)** Transcription factor binding enrichment analysis from genes upregulated in the bulk RNAseq for the indicated conditions. **(e)** Representative examples of gene expression changes across the conditions. Note increased expression of differentiation genes in basal compressed and hypertonic shock conditions. **(f, g)** Schematic of experimental outline (f), representative images and quantification (g) of iPSCs immunostained for Oct4 and Pax6 after compression or hypertonic shock. Note increased differentiation in compressed cells or cells exposed to hypertonic shock (n= 3 independent experiments with >650 nuclei/experiment; ANOVA/Kruskal-Wallis).

To ask if this compression/hyperosmotic shock-mediated induction of differentiation gene expression in basal media conditions was sufficient to influence long-term lineage progression, we induced spontaneous differentiation of iPSCs by leaving them in basal medium lacking pluripotency-maintenance factors (FGF2 and TGF-β1) after 30 min compression or hypertonic shock (Fig. 3f). Strikingly, this transient 30 min compression or hypertonic shock in basal media, which mimic the mechano-osmotic remodeling of the nucleus at the onset of differentiation (see Fig. 1), accelerated ectodermal differentiation observed 96 h later (Fig. 3g). In contrast, and consistent with the transcriptome analyses, transient 30 min compression in the pluripotency medium delayed biochemically-induced differentiation into ectoderm (Supplementary Fig. S6b). Taken together this data indicated that hyperosmotic stress in the absence of pluripotency factors promotes spontanous exit from pluripotency, as evidenced by the differentiation-accelerating effects of hypertonic stress and compression in the basal medium. In contrast, mechano-osmotic chromatin priming in the presence of pluripotency factors reinforces the pre-existing cell state.

### Osmotic pressure controls CBX2 condensation to gate gene repression

We next sought to unravel the distinct mechanisms by which mechanical and osmotic stress gate chromatin priming toward exit or sustained pluripotency maintenance. To compare these two stimuli, we quantified the phosphoproteome of cells subjected to 5 min of hypertonic shock or compression in pluripotency conditions, where the mechanical stress response dominated over the osmotic stress response. Hierarchical clustering of the significantly altered phosphosites revealed clusters either shared by or specific to both stimuli (Fig. 4a). A small group of phosphosites specifically regulated by axial compression were related to chromatin remodeling, and in particular, compression had a strong impact on the phosphorylation of the central component of the NuRD chromatin remodeling complex, CHD4 (Fig. 4b, c). CHD4 controls transcriptional repression of specific genomic loci and is under the control of YAP ^57^. Interestingly, phosphosites upregulated by both compression and hyperosmotic shock were related to chromatin remodeling but also negative regulation of transcription (Cluster 2; Fig. 4b, c). Specifically, phosphorylation of components of the Polycomb repressive complex 1^58^ were upregulated (Fig. 4c). Finally, phosphosites upregulated specifically in response to hypertonic shock were regulators of the actomyosin cytoskeleton and Mitogen-activated protein kinases (MAPK) 1, 3 and 14, corresponding to the activation of ERK1/2 and p38 MAPKs (Fig. 4c). Taken together, these results highlight the distinct (YAP) and overlapping (ERK, PRC1) pathways activated by compression and hyperosmotic shock, and suggest potential regulatory mechanisms of chromatin priming.

**Figure 4:**
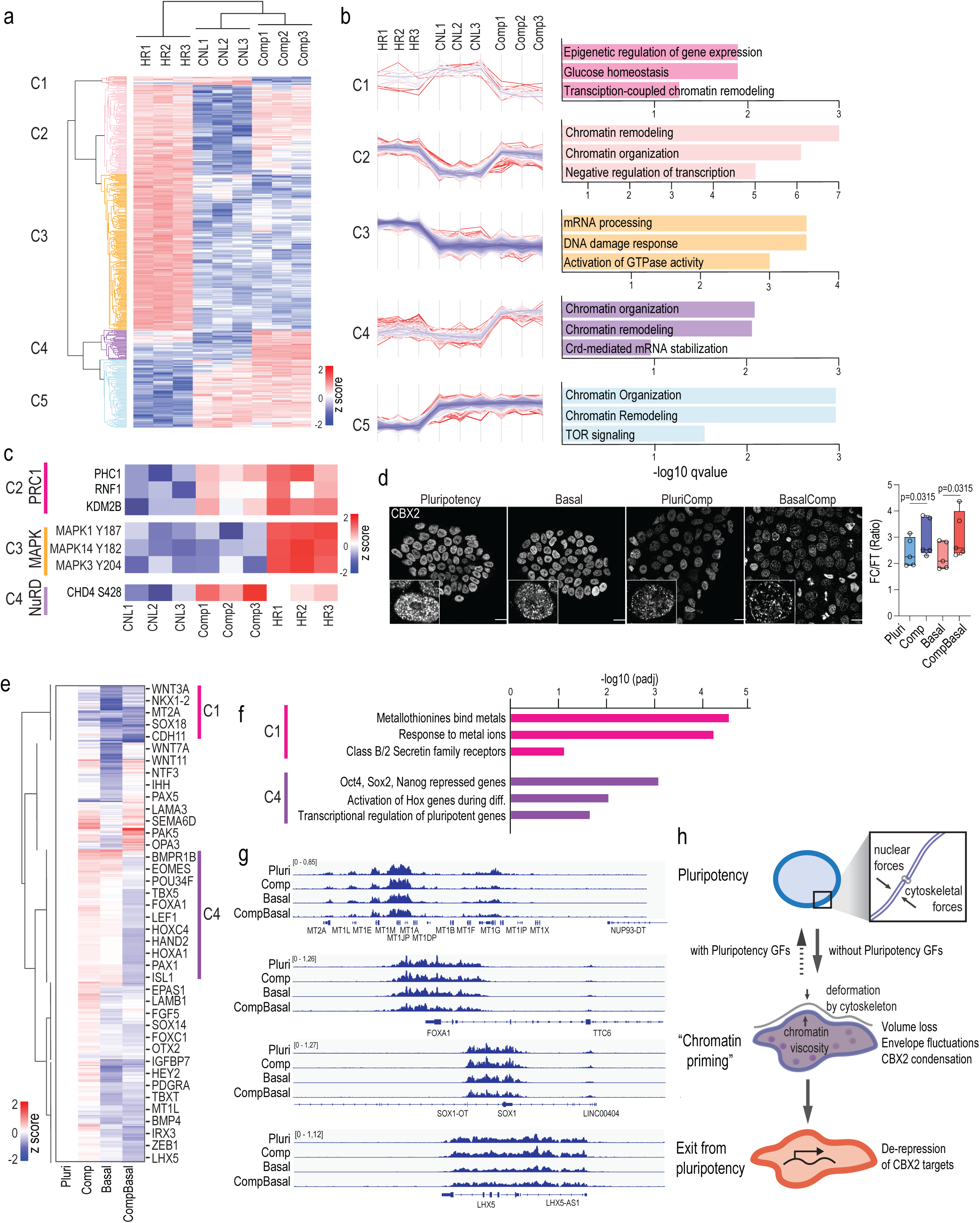
Osmotic pressure controls CBX2 condensation to gate gene repression. **(a)** Heatmap and Euclidian distance dendrogram of differentially abundant phosphosites quantified by mass spectrometry in cells subjected to compression (comp) or hypertonic (HR) stress. **(b)** Distance-based clustering of phosphosites and GO-term analyses show changes specific or common to the specific stresses. **(c)** Example heatmaps of differentially abundant phosphoproteins from (b). **(d)** Representative images and quantification of CBX2 condensation in hiPSCs (Scale bars 20 µm; images representative of n= 5 independent experiments with >20 nuclei/condition/experiment; RM-ANOVA/Holm-Sidak). **(e)** Heatmap and Euclidian distance dendrogram of differential CBX2 occupancy quantified by CUT&Run in cells subjected to removal of pluripotency factors (basal), compression (comp) or compression in basal medium, normalized to pluripotency condition. **(f)** Reactome analysis of genes in clusters 1 and 4 implicate metal-binding genes with reduced CBX2 occupancy in both basal medium and basal medium compression condition whereas pluripotency genes show reduced CBX2 in compression in basal medium. **(g)** Representative tracks of genes with altered CBX2. **(h)** Model of how intranuclear and cytoskeletal forces influence iPSC exit from pluripotency. Under conditions with pluripotency growth factors (GFs), nuclear mechanics are maintained and differentiation is prevented under volumetric stress, restoring pluripotency gene expression. In the absence of pluripotency GFs, osmotic stress leads to nuclear envelope fluctuations and CBX2 condensation, priming chromatin for a cell state transition. This ultimately causes de-repression of CBX2 target genes, facilitating exit from pluripotency.

We first asked if the differential YAP and ERK activation observed by compression versus osmotic stress would explain the enhanced spontaneous differentiation that we observed in the absence of pluripotency factors. To this end we addressed the long-term transcriptional response of withdrawal of pluripotency factors, combined with transient compression and hypertonic shock (30 min followed by 24h recovery) on differentiation gene expression. These experiments confirmed that the osmotic stress response (as indicated by metallothionein gene induction) as well as spontaneous differentiation were most strongly triggered by compression in the basal medium. Inhibition of ERK and YAP did not prevent spontaneous differentiation, but instead enhanced the expression of differentiation genes, and especially in the context of compression in the pluripotency medium (Supplementary Fig. S7a). This effect is consistent with the established role of ERK and YAP in alleviating osmotic and mechanical stress, respectively^59, 60^. Thus, these experiments suggested that ERK and YAP activation function to mitigate mechano-osmotic stress and thus their downregulation subsequently accelerates exit from pluripotency.

As ERK and YAP signaling were not sufficient to explain why osmotic stress triggers chromatin priming for pluripotency exit, we returned to examine chromatin regulators whose phosphorylation was altered in these conditions. Focusing on protein complexes that were co-regulated in the phosphoproteome, we turned to examine the PRC1 pathway, supported by the enrichment of target genes of its central component CBX2 in the 24 h recovered transcriptome in response to transient axial compression in basal medium (Fig. 3d). As serine phosphorylation has been associated with increased propensity to form condensates ^61^ and macromolecule concentration is likely to be altered by the observed changes in nuclear volume and nucleoplasm crowding/viscosity^62^, we analyzed CBX2 condensation in response to axial compression as well as upon removal of pluripotency factors. High resolution imaging of CBX2 immunostaining showed that both removal of pluripotency factors as well as axial compression increased CBX2 condensation (Fig. 4d). Interestingly, these condensates frequently localized at the nuclear periphery together with lamina-associated heterochromatin, potentially sequestering CBX2/PRC1 away from some of its target genes (Supplementary Fig. S7b). To test this hypothesis and the long-term effects of mechano-osmotic CBX2 condensation, we profiled CBX2 occupancy genome-wide 24h after removal of pluripotency-promoting factors with or without an initial 30 min pulse of compression. As expected, CBX2 was found occupying differentiation genes as well as known PRC1/2 targets ^63^ (Supplementary Fig. S7c). Strikingly, while removal of pluripotency factors led to reduced CBX2 occupancy at key differentiation genes, the reduction was strongest when a pulse of axial compression was applied (Fig. 4e, f). Consistent with the reversibility of chromatin changes observed in the ATACseq, axial compression in pluripotency-promoting conditions had only a minor effect on CBX2 occupancy long term (Fig. 4e, f). In contrast, upon compression in the basal medium CBX2 occupancy was substantially reduced at genes repressed by pluripotency factors as well as HOX genes that are involved in differentiation (Fig. 4e-g). Interestingly, the osmoresponsive metallothionein gene cluster was occupied by CBX2, and this occupancy was also moderately reduced by compression in the pluripotency medium, but more strongly affected in the basal medium or upon axial compression in the basal medium (Fig. 4e-g). Collectively these experiments demonstrated that mechano-osmotic forces trigger redistribution of CBX2 condensates and genome-wide occupancy, leading to de-repression of genes that are involved in buffering against osmotic stresses as well as differentiation genes.

## Discussion

The highly regulated three-dimensional organization of genomes arises from interactions across scales, starting from DNA loops to chromatin domains, and higher-order compartments. Despite the dynamics and stochasticity of transcription and variability of chromatin architecture at the single-cell level, these processes are tightly regulated and gate cell fate transitions. The dynamic, probabilistic properties of genome organization have raised questions on how cell states are generated and maintained on the level of single cells and how a population of cells are able to coordinately transition between cell states ^64, 65^. Here we demonstrate that rapid mechano-osmotic remodeling of chromatin architecture and mechanics influences lineage transition in pluripotent cells. Our conclusions are based on comprehensive mechanical and genomic analyses of nuclear states. Upon removal of pluripotency-promoting factors, we observe that nuclear deformation and fluctuations are enhanced, driven by a combination of cytoskeletal confinement and nuclear dynamics. The resulting changes in osmotic stress and nucleoplasm rheology promote chromatin remodeling to accelerate lineage transition. These growth factor-controlled mechanisms can be mimicked and further amplified by inducing nuclear deformation through mechanical confinement or osmotic stress.

We show that in addition to controlling specific gene expression programs on longer time scales, removal of pluripotency growth factors mediates rapid, minute-scale changes in the mechano-osmotic state of the nucleus, characterized by increased energy-dependent nuclear fluctuations and decreased nuclear volume, leading to increased nucleoplasm viscosity and macromolecular crowding. These effects can be enhanced by mechanical flattening of the nucleus. We conclude that these changes in nuclear fluctuations result from an altered force balance between chromatin and the cytoskeleton, as has been described before in other experimental scenarios ^36^. On the level of chromatin, particularly the osmotic component of nuclear deformation enhances accessibility of differentiation genes, providing a state of chromatin priming observed across the entire cell population. We propose that while this transient and reversible chromatin remodeling is not sufficient to facilitate cell fate conversion, it lowers the energy barrier for a signaling factor-driven cell state change and possibly synchronizes chromatin state of a heterogenous cell population to be equally receptive for the specific signaling factors (Fig. 4h).

Our data further indicates that nuclear deformation through confinement has two components: a mechanical component that triggers specific enhancer changes and transcriptional responses mediated most likely by mechanosensitive transcription factors such as YAP. The second component is osmotic stress arising from nuclear volume loss, triggering a transcriptional reset and chromatin remodeling. Our work identifies the PRC1/CBX2 axis as a mechano-osmotically sensitive chromatin regulator. This is consistent with previous work implicating components of the CBX2-PRC1 complex form condensates that co-localize with chromatin and genes relevant for their function ^66–68^, as well as the importance of condensate formation as a mechanism to buffer cells from osmotic pressure and heat stress ^61^. Interestingly, we find that metallothionein genes that are involved in intracellular metal chelators for metal detoxification and scavenging in response to various cellular stresses ^56^ are highly sensitive to mechanical stress and growth factor removal but are also targets of CBX2 repression, consistent with previous work ^63^. Due to the negative charge of DNA, it has high binding propensity with metal ions that in turn are capable of modulating its conformational state as well as interactions with transcription factors ^69^. Interestingly, transport of zinc into the nucleus has been shown to coincide with spontaneous differentiation of human ESC ^70^. This suggests potential existence of a transcriptional feedback loop where a decrease in chromatin repression and increased remodeling is associated with increased transcription of metal detoxifying genes to prevent their impact on DNA. Interestingly, a number of studies identify metallothionein genes and heavy metals in cell state transitions ^8, 70–72^, further implicating mechano-osmotic remodeling as a potential mechanism in regulation of transition dynamics in various scenarios. Future work is required to define how growth factor signaling precisely integrates into this transcriptional regulatory loop.

While a large number of studies have implicated both growth factor signaling and mechanical forces in regulating gene expression, the epigenome, and cell states ^19, 73–75^, it has remained unclear how forces interplay with biochemical signals to determine transcriptional outcomes as well as the physiological relevance of such signal integration. This study provides a foundation for understanding the role of mechanochemical feedback loops, where biochemical signals are capable of altering mechanical properties of the nucleus to change chromatin states, while extrinsic forces have the capacity to alter the way by which biochemical signals are interpreted by cells to impact the kinetics of cellular behaviors.

## Methods

### Cell Culture

Human iPSCs were purchased from Allen Institute (Allen Cell Collection). Sox2-GFP tagged cells were used for experiments unless indicated otherwise. Cells were cultured on Matrigel-coated plates in mTeSR (StemCell technologies) medium and placed in E6 basal medium (Gibco) directly at the onset of experiments wherever indicated. TGF-β1 (5 ng/mL) and bFGF (100 ng/mL) were purchased from StemCell Technologies and were supplemented to basal media where indicated.

### iPSC trilineage differentiation

Sox2-GFP iPSCs were differentiated into mesoderm, endoderm and ectoderm lineages according to the STEMdiff Trilineage Differentiation protocol (StemCell Technologies, #05230). Briefly, cells were seeded on Matrigel-coated plates 5×10^4^ cells/cm^2^ for mesoderm induction or 2×10^5^ cells/cm^2^ for endoderm and ectoderm induction, and were treated the following day with their respective STEMdiff media until analyzed. At 0, 8, 24 and 48 hours post differentiation induction, cells were either imaged live or fixed and stained as detailed below.

### Human embryos and ethical issues

Collection and experiments on human embryos were approved by the Helsinki University Hospital Ethics Committee (diary number HUS/1069/2016). Research permission was approved by the Helsinki University Hospital Research Committee. Human surplus blastocysts were donated by couples that had been treated for infertility at the Helsinki University Hospital Reproductive Medicine unit. The samples were donated with an informed consent, and patients understood that donating embryos is voluntary. Day 5 and day 6 blastocysts (4AA, 4AB or 4BA) vitrified with either Kitazato Vitrification Media (VT601, Kitazato) or Cryotch Vitrification Solutions (110, Cryotech) were warmed with Kitazato Thawing Media (VT602, Kitazato). Cryotec strips (Cryotech) were quickly immersed into the TS (+37), incubation 1 min. The blastocysts were gently aspirated and transferred to the bottom of the DS (RT). After a 3 min incubation in the DS, the blastocysts were gently transferred to the bottom of the WS1 and incubated for 5 min. Subsequently the blastocysts were transferred to the top of the WS2 and after sinking to the bottom, the blastocysts were again transferred to the top of WS2 and after sinking, they were transferred to culture media (GTL, Vitrolife) and cultured in a GERI incubator for 15-16 h.

### Immunofluorescence and confocal microscopy

Human blastocyst stage embryos were fixed in 3.8% paraformaldehyde (PFA) for 15 minutes at room temperature, washed three times in washing buffer (0.1% Tween20-Dulbecco’s Phosphate Buffered Saline (DPBS)) and permeabilized in 0.5% Triton-X100 -DPBS for 15 minutes at room temperature. After washing the embryos in washing buffer, unspecific binding of primary antibodies was blocked by incubating the embryos in Ultra Vision Protein block (Thermo Fisher Scientific) for 10 minutes at room temperature. The embryos were then incubated in primary antibodies at 4°C for overnight. After washing the embryos three times in washing buffer, the embryos were incubated with secondary antibodies (each diluted 1:500 in washing buffer) for 2 h at room temperature. The embryos were washed three times in washing buffer and DNA was counterstained with DAPI (diluted 1:500 in washing buffer). The embryos were imaged in DPBS either on optical grade plastic μ slide 8 well chambers (Ibidi) or on glass bottom dishes (Mattek).

iPSCs were fixed in 4% PFA, permeabilized with 0.3% Triton X-100 in PBS, and blocked in 5% bovine serum albumin (BSA). Samples were subsequently incubated overnight in primary antibody in 1% BSA/0.3% Triton X-100/PBS, followed by washing in PBS and incubation in secondary antibody in 1% BSA/0.3% Triton X-100/PBS. Finally, cells were imaged directly after staining in PBS or subsequently mounted in Elvanol.

The following primary antibodies were used: OCT3/4 (Santa-Cruz Biotechnology, sc-5279; 1:1000), Brachyury (R&D Systems, AF2085; 1:1000), GATA6 (AF1700, RnD Systems; 1:200), NANOG (D73G4, Cell Signaling Technologies; 1:200), LAMINB1 (66095-1-Ig, Proteintech; 1:200), Pax6 (Invitrogen, #42-6600; 1:1000), SOX1 (R&D Systems, AF3369; 1:200), SOX7 (R&D Systems, AF1924; 1:1000), YAP1 (Santa Cruz sc-101199; 1:200), CBX2 (Thermo Fisher PA-582812; 1:800). Alexa Fluor 488, 568, 594 and 647 conjugated antibodies (all from Invitrogen) were used as secondary antibodies at 1:500 dilution.

Fluorescence images were collected by laser scanning confocal microscopy (LSM980; Zeiss) with Zeiss ZEN Software (Zeiss ZEN version 3.7), or with Andor Dragonfly 505 high-speed spinning disk confocal microscope (Oxford Instruments) equipped with 488 nm and 546 nm lasers, and an Andor Zyla 4.2 sCMOS camera using 40x, 63x or 100x immersion objectives. Images were acquired at room temperature using sequential scanning of frames of 1 µm thick confocal planes (pinhole 1). Images were collected with the same settings for all samples within an experiment.

### Segmentation and image analysis

#### Nuclear volume quantification

LaminB1-RFP expressing hiPSCs were plated on glass bottomed imaging disheds coated with Matrigel. Cell colonies were live imaged for 10 min (growth factor experiment) or 40 min (cell compression experiments) acquiring full z-stack from same colony for each timepoint. Images were acquired using Andor Dragonfly spinning disk confocal microscope in environmental chamber (5% CO2 and + 37°C. The volumes were quantified using ImageJ ^76^. 4D movies were filtered using 3D median filter and subsequently bleach corrected using simple ratio correction. Individual nuclei were identified manually and marked with an oval selection stored to ROI manager. Seeds were then expanded using the Limeseg-plugin ^77^. The segmentation was manually supervised to ensure complete segmentation. In the case of human embryos, the segmentation was performed similarly with following modifications: Fixed immune stained embryos were imaged using 40x oil immersion objective, and cells were identified as inner cell mass based on Nanog expression. ICM cells with high expression of Nanog or Gata6 were then manually identified and segmented based on LaminB1 signal. For volume calculation in 3-lineage differentiation experiments Sox2-GFP-expressing hiPSCs cells were plated on glass-bottom 35 mm gridded bottom dish (Ibidi) to image the same colonies of cells across multiple timepoints. A complete z stack was imaged using a Nikon eclipse Ti2 inverted microscope mounted with a CSU-W1 spinning disk microscope (60X Oil Immersion Lens; NA = 1.49) immediately prior to starting the differentiation protocol. The same colonies were imaged 8 hours, 24 hours, and 48 hours after the addition of differentiation media. 3D segmentation of nuclei (outlined by Sox2-GFP expression) was performed using a custom Cellpose (version 2.2.2) model ^78, 79^ to generate 3D masks. 3D masks were converted to 3D ROIs using the 3D Suite plugin ^80^, after which nuclear volumes was quantified from 3D ROIs using the same plugin.

#### CBX2 cluster analysis

CBX2 immunostained samples were imaged with LSM980 with 63x oil immersion objective using Airy Scan imaging mode and subsequent deconvolution. Individual cell nuclei were segmented from the images using custom Cellpose (version 2.2.2) model ^78, 79^. CBX2 clusters were detected using cellpose and custom trained model recognizing small cluster from the overall signal. The mean intensity of CBX2 was measured from background subtracted images from the cluster area and non-clustered area. Subsequently cluster enrichment (mean intensity cluster: mean intensity nucleus) was calculated for each cell. The same segmented areas were used to calculate the relative amount of DNA at the CBX2 cluster using the Dapi staining. The mean intensities were measured with ImageJ^76^ and morpholib - plugin ^81^.

For localization at the nuclear periphery nuclei were segmented from high resolution microscopy images using a custom Cellpose (version 2.2.2) model. Masks were dilated isotropically 400nm. The dilated masks were then eroded 20 times for 150 nm in order to generate thin consecutive masks using ImageJ. The thin masks were used to measure spatially resolved mean intensity of CBX2 and Dapi from nuclear periphery from background subtracted images. Each measurement was then divided by mean intensity of either CBX2 or Dapi from the whole nuclear area to calculate spatially resolved peripheral enrichment.

#### Quantification of nuclear intensities of transcription factors

Mean intensities were measured from images using a nuclear mask generated from DAPI or Sox2-GFP using ImageJ ^76^. N/C ratio for YAP staining was calculated for each cell by dividing nuclear mean intensity with cytoplasmic mean intensity.

### Mechano-osmotic perturbations

Compression was performed with a modified version of a previously published cell confiner system^82^. Briefly, suction cup-bound coverslips were driven on 2D iPS cell colonies using a controlled pressure pump (Elveflow). Compression height was controlled by 5 µm polystyrene bead spacers. Compression was maintained on the nuclei for indicated times in the presence/absence of chemical treatments or media conditions, as indicated, and cells were harvested or fixated in 4% PFA directly at the end of the compression period or at a recovery time point indicated.

For the western blotting, phosphoproteomics and sequencing experiments cells were axially compressed using polystyrene block custom manufacture to fit 6 cm or 10 cm cell culture dishes. Shortly, the polystyrene weight was heated to +37C degrees in cell culture incubator prior to the experiment. The cells were once rinsed with warm PBS and media was changes to minimal amount to cover the culture plate. The block was then place on top of the cells starting from one corner and inducing spread of homogenous layer of culture medium between the block and the cells.

Hyperosmotic shock was induced using solution of 1M Sucrose. Pre-heated (+37 C) sucrose solution was added to the cell culture medium to achieve 50:50 mixture. Hypo-osmotic shock was induced using pre-heated mQH_2_0 that was added to the medium to achieve a final 50:50 mixture.

### Generation of Halotag-YAP1 knockin iPSCs

0.6pmol (∼1.6ug) pUC57-Halotag-N-YAP1 plasmid (custom made by Genscript, containing 400bp/ea homology arms flanking YAP1 (NM_001130145.3) start codon and Halotag coding sequence followed by GGSGGS linker) was mixed with 12.2pmol Alt-R HiFiCas9 v3 (IDT #1081061) and 17.5pmol sgRNA (Synthego, 5’-CGGCTGCTGCCCGGGATCCA) to transfect 226,000 WTC-dCas9-TagBFP-KRAB iPSCs (Coriell # AICS-0090-391) using Nucleofector 4D 16-well cuvette strip, 20uL P3 Primary Cell Nucleofector Solution (Lonza #V4XP-3032), and program CA-137. After nucleofection, the iPSCs were cultured in one well of rhLaminin-521 (Thermo #A29249) coated 24-well plate with 0.6mL StemFlex medium (Thermo #A3349401) plus 1uM HDR enhancer v2 (IDT #10007921) and 1x RevitaCell (Thermo #A2644501) in 32°C incubator overnight. Then the medium was changed to StemFlex plus 0.5x RevitaCell. 3 days after nucleofection, the iPSCs were moved to 37°C incubator with fresh StemFlex medium change every other day. 16 days after nucleofection, the iPSCs were incubated with 1µM Halotag Oregon Green ligand (Promega # G2802) following manufacturer’s rapid labelling protocol, and the Oregon Green positive cells were sorted into 1 cell/96-well in Matrigel-coated 96-well plate with 1x CloneR2 (Stem Cell Technologies, # 100-0691) and 100µL/well StemFlex using BD FASCMelody cell sorter.

To confirm Halotag-YAP1 knockin iPSC clones only have 1 or 2 copies of Halotag knockin allele and no vector integration, ddPCR (Bio-rad QX200) was used with Halotag primer (5’-GCTCCGCTCTGGGTTTC, 5’-GGGCGGATGAACTCCATAAA) and probe (/56-FAM/TCCAGAGCG/ZEN/CGTCAAAGGTATTGC/3IABkFQ/), AmpR primer (5’-TTTCCGTGTCGCCCTTATTCC, 5’-ATGTAACCCACTCGTGCACCC) and probe (/56-FAM/TGGGTGAGC/ZEN/AAAAACAGGAAGGC/3IABkFQ/), and reference hRPP30 (5’-GGCCATCAGAAGGAGATGAAG, 5’-AAGGGAGTGCTGACAGAGA) and probe (/5HEX/AGAAAGCCA/ZEN/AGTGTGAGGGCTGAA/31ABkFQ/). Clones with 2 copies of Halotag and zero copy of AmpR were selected as homozygous Halotag-YAP1 knockin clones. The knockin alleles in these clones were further confirmed by genomic PCR of a 2.5kb fragment using primers (5’-GCGTTTGAGGCGAGTTTCTG, 5’-AAGCGTTTCAGCCGACTGTA) followed by Sanger sequencing using primers (5’-GCGTTTGAGGCGAGTTTCTG, 5’-CTCCAGTTCCACTTCGCCTC).

### Nuclear envelope fluctuation analyses

For measuring NE fluctuations for approximating apparent NE tension, RFP-LMNB1 iPSC colonies were imaged using a high frame rate acquisition mode (150 ms /frame) on a spinning disc microscope (Andor Dragonfly) for 5 min. Images were then corrected for bleaching (Bleach Correction function of ImageJ) and linear rotational drift (Stackreg; ImageJ). Following corrections, the position of nuclear edge was recorded as a function of time at different positions along the NE by drawing a line perpendicular to NE at multiple locations per nucleus. NE fluctuations were calculated by measuring the standard deviation of the position of the NE from its mean position as described previously^37^. ATP was depleted using Actinomycin D (10 µM) and 2-desoxyglucose (5mM) and the cytoskeleton was disrupted with Cytochalasin D (200 nM) and Nocadozole (400 nM) where indicated. For live imaging of the actin cytoskeleton the cells were stained using SPY650-FastAct (SpiroChrome).

### Atomic force microscopy (AFM)

AFM force spectroscopy measurements were performed on hiPS cell nuclei using JPK NanoWizard 2 (Bruker Nano) atomic force microscope mounted on a Nikon Eclipse Ti inverted microscope and operated with JPK SPM Control Software v5. Glass-bottom cell culture dishes were mounted on the AFM directly after switching cell culture media and measurements were performed at 37℃ within 20 min. Triangular non-conductive silicon nitride cantilevers (MLCT, Bruker) with nominal spring constant of 0.01Nm^-1^ were used for force spectroscopy measurements of the apical surface of cells directly over the nucleus, where the tip was directly aligned using brightfield view. For all indentations, forces of up to 3nN were applied and the velocity of indentation was kept constant at 2 µm s^-1^ ensuring an indentation dept of approximately 500 nm. A force spectroscopy map of 4 µm^2^ with resolution 2×2 px was used to perform technical replicates of nuclei indentation, and all valid curves were analyzed. Prior to fitting the Hertz model to obtain Young’s Modulus (Poisson’s ratio of 0.5), the offset was removed from the baseline signal, the contact point was identified, and cantilever bending was subtracted from each force curve. All analysis was performed with JPK Data Processing Software (Bruker).

### NucGEMS

hIPSC were transfected in suspension during plating using X-tremeGENE (Sigma) according to manufacturer’s instructions. Colonies were imaged 36h post transduction on a Nikon Eclipse Ti Eclipse microscope mounted with Yokogawa CSU-W1 spinning disk unit using 405 and 488 laser and 63X CFI Apo 60x/N.A-1.49/.12 TIRF objective and ET525/36m emission filter (Chroma). Images were acquired using 80% power from single focal plane at 100ms intervals, 1×1 binning, 16-bit pixel depth. High-Throughput Image Processing Software platform ^83^ was used to track particles and custom python pipelines were used to calculate aggregate mean square displacement, effective diffusivity and the diffusive exponent ^84^. Single particle tracks were generated with Napari GEMspa plug-in ^85^.

### Multiome sequencing and analysis

Cells were compressed as described above after which single nuclei were harvested using Nuclei buffer and following manufacturer instructions (Chromium Next GEM Single Cell Multiome ATAC + Gene Expression kit). Single-nuclei encapsulation and library preparation were performed according to the manufacturer’s protocol. Two biological replicates were prepared for each condition and sequenced using the Chromium Single Cell Multiome platform (10x Genomics). All replicates were quality controlled and analyzed separately to ensure reproducibility after which all conditions were pooled and analyzed together, treating each cell in the total pool as a biological replicate. Initial transcript count and peak accessibility matrices were obtained with Cell Ranger Arc (v.1.1.2).

Raw counts were subsequently imported into Python (3.8) as AnnData (0.8.0) objects. Cells with more than 25% mitochondrial RNA content were removed. Doublet prediction on scRNA-seq data was performed using scrublet (0.2.3). Raw scRNA-seq counts were normalized using scran (1.22.1) with Leiden clustering input at resolution 0.5. scRNA-seq and scATAC-seq data were integrated as MuData objects (0.2.2) using Muon (0.1.2). Raw scATAC-seq counts were filtered for noise, and subsequently TF-IDF transformed using muon.atac.pp.tfidf with scale factor 10,000. scRNA-seq data was further processed using scanpy (1.8.2): for 2D embedding, the expression matrix was subset to the 2,000 most highly variable genes (sc.pp.highly_variable_genes, flavor “seurat”). The top 50 principal components (PCs) were calculated, and served as basis for k-nearest neighbor calculation (sc.pp.neighbors, n_neighbors=30), which were used as input for UMAP (<https://doi.org/10.48550/arXiv.1802.03426>) layout (sc.tl.umap, min_dist=0.3). scATAC-seq data dimensionality was reduced to 50 components using muon.atac.tl.lsi, and embedded in a 2D UMAP (sc.tl.umap) on the basis of k-nearest neighbor calculation (sc.pp.neighbors, n_neighbors=20).

Gene signatures for cell cycle ^86^ apoptosis (BCL2L1, CASP9, CYCS, IL1A, PIK3CG, TNFRSF10D, FADD, BIRC3, FAS) and necrosis (BIRC3, FAS, DNM1L, GSDME, IPMK, MLKL, RBCK1, TICAM1, YBX3) were scored using the scanpy “sc.tl.score_genes” function. Pychromvar (0.0.4) was used to interrogate ATAC-seq transcription-factor accessibility. Pseudobulk differential expression analysis was performed with pyDESeq2 (0.4.8). Active enhancers and gene regulatory networks were inferred from the multiome data by linking peaks to genes using SCENIC+ ^50^ (pyscenic 0.11.2). SCENIC+ was performed by following steps outlined in the documentation. Briefly, topic modeling was performed with 16 topics after evaluating different LDA models, and binarized using the Otsu method. Condition annotations were added to the metadata alongside topics. Accessibility matrices were imputed and normalized, and highly variable features and condition-specific accessibility were calculated. Custom motif and ranking databases were created using “create_cisTarget_databases” based on the observed scATAC-seq peaks. Motifs were obtained from the SCENIC+ “v10nr_clust_public” collection. After running SCENIC+, eRegulons underwent standard filtering and scoring.

### H3K27ac Cut & Tag and analysis

Cut&Tag was performed using a Complete CUT&Tag-IT Assay Kit (Active Motif) according to manufacturer’s instructions. Conditions were plated in triplicate and subject to 30 min treatments as indicated. Cells were collected from 10 cm dishes by brief incubation at 37℃ in 0.05M EDTA in HBSS to promote detachment followed by cell scraping. Briefly, cells were attached to Concanavalin-A-conjugated magnetic beads (ConA beads, Polysciences) before overnight binding of primary antibody H3K27ac (Abcam, ab4729). Samples were then incubated with secondary antibody and CUT&Tag-ITAssembled pA-Tn5 Transposomes, after which tagmented DNA was purified. This DNA was PCR amplified using a unique combination of i5/i7 indexing primers for each sample, cleaned using SPRI beads and pooled into an equimolar library for sequencing on an Illumina HiSeq4000 sequencing platform.

Unmapped paired-end reads were trimmed to remove adapters and poor-quality sequences using fastp v0.23.2 (--detect_adapter_for_pe). Paired-end reads were mapped to GRCh38 and Drosophila melanogaster (GCF_000001215.4_release_6_plus_iso1_mt) reference genomes using bwa-mem2 v.2.2.1 with default settings. To define active enhancers and promoters, we performed peak calling using the callpeak function (“-f BAMPE -g hs -q 1e-5 --keep-dup all –nomodel”) from MACS2 (2.2.7.1) ^87^. PCR duplicates were removed using sambamba (v 1.0.1) ^88^ and the IgG negative control from the CBX2 Cut & Run experiment was used for estimating background. The final set of H3K27ac-enriched genomic regions included a union of peaks from all experimental conditions that were detected in at least two biological replicates, with peaks closer than 1 kb merged together, and peaks overlapping with the ENCODE blacklisted genomic regions discarded ^89^. All H3K27ac peaks were further classified into active promoters (n = 10,728) and putative active enhancers (n = 15,531) based on the overlap with TSS ± 1 kb regions of hg38 UCSC refGene and protein-coding GENCODE v38 genes. To link putative active enhancers with their potential target genes, we used the closest function from BEDTools (v2.31.1) ^90^ to find the nearest UCSC refGene gene (TSS ± 1 kb region).

Following PCR duplicate removal, biological replicates (2 replicates for Pluri and hyper, 3 for all others) from the same condition were spike-in normalized and merged together as described previously ^91^, and then used to generate coverage tracks using bamCoverage from deepTools (v 3.5.4) ^92^(“-bs 1 – ignoreDuplicates”). BigwigCompare from deepTools (“--skipZeroOverZero --operation log2 -- fixedStep -bs 50”) was used to generate the differential H3K27ac enrichment tracks. To perform metaplot and heatmap analysis of the mean read density at regions of interest, we used computeMatrix and plotProfile/plotHeatmap from the deepTools suite.

Significant changes (p-adj < 0.05, |fold change| > 1.5) in H3K27ac enrichment at active enhancers following different treatments were identified using DESeq2 R package (v 1.42.1) ^93^as described previously ^94^. Briefly, multiBamSummary from deepTools (--outRawCounts) was used to obtain counts of hg38 mapped reads at target regions of interest across different conditions, following PCR duplicate removal. Read counts from the spike-in dm6 genome at dm6 refGene genes were used to calculate size factors for spike-in calibrated DESeq2 analysis ^91^. To visualize an overlap of active enhancers decommissioned following compression (a union of peaks showing a significant loss of H3K27ac in the compressed pluripotency and basal conditions, n = 3,694) and hyperosmotic shock (n = 5,496), we used UpSetR (v 1.4.9) and VennDiagram (v 1.7.3) R packages. GO term analyses were carried out using Enrichr ^95^.

### Nascent RNA sequencing (TT-seq) and analysis

TTseq was performed essentially as described previously ^52^. Briefly, cells were labeled with 500 µM 4-thiouridine (4sU) for the last 10 min of the indicated 30 min treatments, after which cells were harvested and lysed in Trizol. ERCC spike-ins (00043, 00170, 00136) were added to lysates after which RNA was isolated, fragmented and biotinylated with EZ-link HPDP-biotin (Thermo Fischer. Biotinylated nascent RNAs were purified with streptavidin-conjugated magnetic beads (µMACS; Miltenyi) and libraries of total and biotinylated RNA were prepared with Illumina TruSeq total RNA kit. Libraries were quantified using the KAPA Library Quantification Kit and sequenced with Illumina NextSeq 500 using the High Output Kit v2.5 (150 cycles, Illumina) for 2 × 75 bp paired-end reads. Trimmed reads were aligned to the human genome assembly (hg38) using STAR2.4.2 ^96^. For coverage profiles and visualization, reads were uniquely mapped, de-stranded, antisense corrected, and normalized with size factors calculated from DESeq2 ^93^.

To quantify changes in the nascent RNA levels from TT-seq, we used a custom script to perform spike-in calibrated differential analysis as described previously ^93^, with spike-in read counts provided for estimating DESeq2 size factors. Raw read counts following PCR removal were obtained for a set of hg38 MANE RefSeq genes (v1.3)^10^ using the featureCounts function (“isPairedEnd = TRUE, countReadPairs = TRUE, useMetaFeatures = TRUE, strandSpecific = 1”) from the Rsubread package (v 2.16.1). Log2 fold changes following shrinkage using the original DESeq2 estimator^7^ were visualized using custom R scripts and ggplot2.

### Bulk RNA sequencing and analysis

Total RNA was isolated using the NucleoSpin RNA Plus kit (Macherey&Nagel). After quantification and quality control using Agilent 2200 TapeStation, total RNA amounts were adjusted and libraries were prepared using the TruSeq Stranded Total RNA kit with Ribo-zero gold rRNA depletion (Illumina). RNA sequencing was carried out on Illumina NextSeq 500 using the High Output Kit v2.5 (150 cycles, Illumina) for 2 × 75 bp paired-end reads.

Raw FASTQ files were adapter-trimmed and quality-filtered using fastp (0.23.2) using default settings and the “—detect_adapter_for_pe” flag. Filtered reads were mapped to the GRCh38 reference genome merged with ERCC92 spike-in sequence reference using STAR (2.7.10a) (-- outSAMstrandField intronMotif --outFilterIntronMotifs RemoveNoncanonical -- outFilterMultimapNmax 1 --winAnchorMultimapNmax 50 --outFilterType BySJout -- alignSJoverhangMin 8 --alignSJDBoverhangMin 1 --outFilterMismatchNmax 999 -- outFilterMismatchNoverReadLmax 0.04 --alignIntronMin 20 --alignIntronMax 1500000 -- alignMatesGapMax 1500000 --chimSegmentMin 15 --chimOutType WithinBAM -- outSAMattributes All). Gene expression counts were generated from mapped reads using Gencode v35 and ERCC92 annotations with htseq-count (2.0.3).

PCA and differential gene expression analysis were performed in R (4.1.2) using DESeq2 (1.34.0) under with the assumption of negative binomial distribution of read counts (Wald test). GO term and transcription factor binding analyses were carried out using Enrichr/ChEA3 ^95^.

### Phosphoproteomics and analysis

hIPSCs in pluripotency media were subjected to compression or hypertonic shock for 5 mins after which cell lysates were collected in 6 M guanidinium chloride buffer supplemented with 5 mM Tris(2-carboxyethyl)phosphine, 10 mM chloroacetamide in 100mM Tris-HCl. Samples were then boiled at 95°C for 10 min, sonicated at high performance for 10 cycles (30 s on/off) using Bioruptor Plus Ultrasonicator (Diagenode), and spun down for 20 minutes at room temperature at 20000 g. Supernatants were then Trypsin-Gold digested (Promega Corp., V5280) overnight at 37°C. Digested samples were acidified and peptides were cleaned with custom-packed C18-SD Stage Tips. Eluted peptides were vacuum dried at 30°C and phosphopeptides were enriched using 3 mg/200 mL Titansphere Phos-TiO kit (GL Sciences) according to manu-facturer instructions. After elution phosphopeptides were vacuum dehydrated for 2 hours at 30°C, cleaned with custom-packed C18-SD. Ultimate 3000 ultra-high performance liquid chromatography (UHPLC) in conjunction with high-pH reversed-phase chromatography was used to separate and fractionate a total of 1 mg of tryptic peptides, from which 8 fractions were collected.

All samples were analyzed on a Q Exactive Plus Orbitrap mass spectrometer that was coupled to an EASY nLC (both Thermo Scientific). Peptides were loaded with solvent A (0.1% formic acid in water) onto an in-house packed analytical column (50 cm, 75 µm inner diameter, filled with 2.7 µm Poroshell EC120 C18, Agilent). Peptides were chromatographically separated at a constant flow rate of 250 nL/min using the following gradient: 3-5% solvent B (0.1% formic acid in 80 % acetonitrile) within 1.0 min, 5-30% solvent B within 121.0 min, 30-40% solvent B within 19.0 min, 40-95% solvent B within 1.0 min, followed by washing and column equilibration. The mass spectrometer was operated in data-dependent acquisition mode. The MS1 survey scan was acquired from 300-1750 m/z at a resolution of 70,000. The top 10 most abundant peptides were isolated within a 1.8 Th window and subjected to HCD fragmentation at a normalized collision energy of 27%. The AGC target was set to 5e5 charges, allowing a maximum injection time of 55 ms. Product ions were detected in the Orbitrap at a resolution of 17,500. Precursors were dynamically excluded for 25.0 s. All mass spectrometric raw data were processed with MaxQuant (version 2.4.0) ^97^ using default parameters against the Uniprot canonical Human database (UP5640, downloaded 04.01.2023) with the match-between-runs option enabled between replicates and phosphorylation at S, T, and Y defined as variable modifications. Results were loaded into Perseus (version 1.6.15) ^97^. and after removing contaminants and insecure identifications. Remaining identified phosphosites were filtered for data completeness in at least one replicate group and remaining data normalized by median subtraction for each individual samples. After sigma-downshift imputation using standard settings, significantly changed sites were identified using both one-way ANOVA and FDR-controlled T-tests. Afterwards, sites were annotated using PhosphoSitePlus ^98^ information. Based on annotation information, 1D enrichments were performed on site abundance differences. ANOVA-significant site abundances were Z-scored and hierarchically clustered using standard settings. GO term analyses were carried out using Enrichr ^95^.

### Western blotting

Cells were rinsed in PBS, suspended in lysis buffer (50 mM Tris-HCl buffer (pH 8.0), containing 150 mM NaCl, 1% Triton X-100, 0.05% sodium deoxycholate, 10 mM EDTA, protease and phosphatase inhibitors) and cleared by centrifugation. The lysates were then reduced in Laemmli sample buffer at 95°C, separated by polyacrylamide gel electrophoresis in the presence of SDS and transferred onto PVDF membranes. Membranes were blocked with 5% milk powder in Tris-buffered saline containing 0.05% Tween (TBS-Tween) for 1 h at room temperature, after which primary antibodies were added in 5% BSA, TBS-Tween and incubated over night at +4°C. The membranes were subsequently washed in TBS-Tween after which secondary horseradish peroxidase conjugated antibodies (Bio-Rad) were added in 5% milk powder in TBS-Tween and incubated for 30 min room temperature. After extensive washing in TBS-Tween, antibody binding was detected by chemiluminescence (Immobilon Western, Millipore) using the Bio-Rad ChemiDoc Imaging System. The following antibodies were used: LaminB1 (Cell Signaling 9087; 1:1000), p38 MAPK phospho Thr180/Tyr182 (Cell signaling 4511, 1:1000), p38 MAPK (Cell Signaling 9212, 1:1000), p44/42 MAPK phospho-Erk1/2 (Cell Signaling 4376, 1:2000), p44/42 MAPK Erk1/2 (Cell Signaling 4695, 1:1000), RNApol2 PS2 (Abcam ab5095; 1:5000).

### RT-qPCR

Cells were treated as indicated with media conditions, compression, and YAP inhibitor verteporfin (200 nM) and ERK inhibitor (1 µM). RNA was isolated using the NucleoSpin RNA plus Kit (Macherey&Nagel). RNA quality was assessed using the high sensitivity RNA Screen Tape Analysis (TapeStation, Agilent) and subsequently 500 ng of RNA was reverse transcribed applying the SuperScript ™ IV VILO Master Mix (Thermo Fisher Scientific) following the manufacturer’s instructions. ERCC RNA Spike-in Mix (Thermo Fisher Scientific) was added to samples as reference to ensure validity of normalization. PCR was performed with PowerUp SYBR Green Mastermix (Thermo Fisher Scientific) using QuantStudio5 Real-Time PCR System (Thermo Fisher Scientific). Gene expression was quantified using the ΔΔCt method with normalization to LaminB1. All of the primers were designed to span exons. The following primer sequences were used:

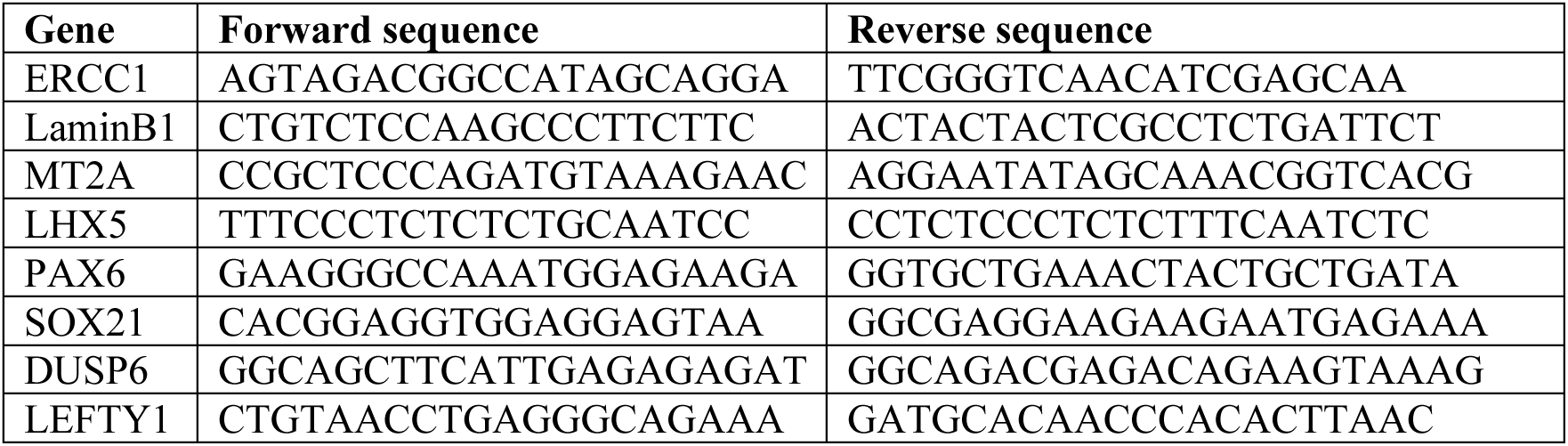

### CBX2 Cut&Run and analyses

Cells were compressed from 30 min in basal or pluripotency medium followed by 24h recovery in the same media condition. Uncompressed cells in the respective media were used as controls (2 biological replicates/condition). CBX2 cut&run was performed using the CUTANA ChIC Cut&Run Kit (EpiCypher) using the manufacturers’ instructions. Briefly, 5000 0000 cells/replicate were harvested and absorbed with activated ConA beads, followed by incubation with 0.5 µg of CBX2 antibody (mAb #25069, Cell signaling) overnight at +4. Chromatin was then digested and released followed by DNA purification and library prep using the CUTANA Cut&Run Library Prep Kit (EpiCypher). After library quantification and quality control using Agilent 2200 TapeStation, DNA sequencing was carried out on Illumina NextSeq 500 using the High Output Kit v2.5 (150 cycles, Illumina) for 2 × 75 bp paired-end reads.

Raw FASTQ files were adapter-trimmed and quality-filtered using fastp (0.23.2) using default settings and the “—detect_adapter_for_pe” flag. Filtered reads were mapped to the GRCh38 and E.coli (ASM886v2) reference genome using bwa-mem2 (2.2.1). Mapped reads were quality-filtered using a cutoff of 3. Scaling-factor normalized tracks were generated according to Zheng et al. (dx.doi.org/10.17504/protocols.io.bjk2kkye). Regions were blacklisted using published CUT&RUN blacklists ^99^.

Coverage profiles for IgG and CBX2 were generated using MACS2 (2.2.9.1). Normalisation to IgG was performed using MACS2 “bdgcmp” and the “qpois” method, and peak calls were obtained using MACS2 “bdgbroadcall” (-c 5 -C 2 -g 1000 -G 2000 --no-trackline). Peaks were filtered for noise using reproducibility (called in > 1 sample of the same condition), minimum peak size (> 1000bp), and minimum mean CBX2 signal in one condition (>0.075).

Aggregate statistics for each sample were calculated on replicate-merged tracks for peaks overlapping known genes (Gencode v44). Mean peak signal was scaled to the pluripotency medium as reference condition, and log2-transformed for symmetry. Hierarchical clustering was performed using Ward method and Euclidian distance metric (scipy 1.11.4), clusters were chosen according to a tree distance cutoff of 7.

### Statistics and reproducibility

Statistical analyses were performed using GraphPad Prism software (GraphPad, version 9) or in R (version 4.2.2). Statistical significance was determined by the specific tests indicated in the corresponding figure legends. Only 2-tailed tests were used. In all cases where a test for normally distributed data was used, normal distribution was confirmed with the Kolmogorov–Smirnov test (α = 0.05). All experiments presented in the manuscript were repeated at least in 3 independent replicates.

### Data availability

All analysis scripts, custom code, and data that support the conclusions are available from the authors on request.

## Acknowledgements

We thank Tom Misteli and Ivan Bedzhov for critical reading of the manuscript, Nada Jabado for advice on CBX2, Matthew Geis and Hunki Lee with help with image segmentation and nucGEMs, Hermann vom Bruch, Claudia Ortmeier and Sandra Heising for expert technical assistance, Fabien Bertillot for advice on image analyses, the Max Planck Institute BioOptics and Sequencing core facilities and the CCR/NCI High-Throughout Imaging Facility (HiTIF), for support with experiments. Human embryos were imaged at the Biomedicum Helsinki Imaging Unit supported by HiLIFE. Multiome sequencing was performed at the Institute for Molecular Medicine Finland FIMM Genomics unit supported by HiLIFE and Biocenter Finland. This work was supported by Instrumentarium Science Foundation and Academy of Finland postdoctoral fellowships (to AS), the Intramural Research Program of the National Institute of Diabetes (NIH) and Digestive and Kidney Diseases (NIDDK; to YAM), the Sigrid Juselius Foundation (to SV and SAW), Helsinki Institute of Life Science (to SV and SAW), and Academy of Finland Research Fellowship (353549 to SV), Academy of Finland Center of Excellence BarrierForce and R’Life Programme consortium NucleoMech (to SAW), German Research Foundation (DFG) FOR 5504 and the Max Planck Society (to SAW).

## Author contributions

KM and AS designed and performed experiments and analyzed data. RS and AC performed and analyzed differentiation experiments. KK performed and supervised sequencing data analysis. JWL performed and analyzed proteomics experiments. AM performed experiments. CMN provided conceptual advice and supervised the proteomics analyses. NAF performed TTseq and H3K27Ac Cut&Tag analyses. LCB supported sequencing experiments. CH-G collected, warmed, and cultured the human embryos. JZ designed and constructed Halo-tagged lines with YAM. AK aided in nucGEM mean square displacement analyses. SN and SV designed and performed all studies on human embryos. YAM supervised the study, performed all sequencing experiments, designed and performed experiments and analyzed data. SAW conceived and supervised the study, designed experiments, analyzed data and wrote the paper. All authors commented and edited the manuscript.

## References

1. Kalkan, T. & Smith, A. Mapping the route from naive pluripotency to lineage specification. Philos Trans R Soc Lond B Biol Sci 369 (2014).

2. Dixon, J.R. et al. Chromatin architecture reorganization during stem cell differentiation. Nature 518, 331–336 (2015).

3. Pelham-Webb, B., Murphy, D. & Apostolou, E. Dynamic 3D Chromatin Reorganization during Establishment and Maintenance of Pluripotency. Stem Cell Reports 15, 1176–1195 (2020).

4. Tyser, R.C.V. et al. Single-cell transcriptomic characterization of a gastrulating human embryo. Nature 600, 285–289 (2021).

5. Lovicu, F.J., McAvoy, J.W. & de Iongh, R.U. Understanding the role of growth factors in embryonic development: insights from the lens. Philos Trans R Soc Lond B Biol Sci 366, 1204–1218 (2011).

6. Pan, G. & Thomson, J.A. Nanog and transcriptional networks in embryonic stem cell pluripotency. Cell Res 17, 42–49 (2007).

7. Brons, I.G. et al. Derivation of pluripotent epiblast stem cells from mammalian embryos. Nature 448, 191–195 (2007).

8. Osnato, A. et al. TGFbeta signalling is required to maintain pluripotency of human naive pluripotent stem cells. Elife 10 (2021).

9. Dattani, A. et al. Naive pluripotent stem cell-based models capture FGF-dependent human hypoblast lineage specification. Cell Stem Cell 31, 1058–1071 e1055 (2024).

10. Linneberg-Agerholm, M. et al. Naive human pluripotent stem cells respond to Wnt, Nodal and LIF signalling to produce expandable naive extra-embryonic endoderm. Development 146 (2019).

11. Simon, C.S. et al. Suppression of ERK signalling promotes pluripotent epiblast in the human blastocyst. bioRxiv, 2024.2002.2001.578414 (2024).

12. Ferrai, C. & Schulte, C. Mechanotransduction in stem cells. Eur J Cell Biol 103, 151417 (2024).

13. Kim, E.J.Y., Korotkevich, E. & Hiiragi, T. Coordination of Cell Polarity, Mechanics and Fate in Tissue Self-organization. Trends Cell Biol 28, 541–550 (2018).

14. Ozguc, O. & Maitre, J.L. Multiscale morphogenesis of the mouse blastocyst by actomyosin contractility. Curr Opin Cell Biol 66, 123–129 (2020).

15. Chen, Q., Shi, J., Tao, Y. & Zernicka-Goetz, M. Tracing the origin of heterogeneity and symmetry breaking in the early mammalian embryo. Nat Commun 9, 1819 (2018).

16. Firmin, J. et al. Mechanics of human embryo compaction. Nature 629, 646–651 (2024).

17. Bertillot, F., Miroshnikova, Y.A. & Wickstrom, S.A. SnapShot: Mechanotransduction in the nucleus. Cell 185, 3638–3638 e3631 (2022).

18. Chugh, M., Munjal, A. & Megason, S.G. Hydrostatic pressure as a driver of cell and tissue morphogenesis. Semin Cell Dev Biol 131, 134–145 (2022).

19. Dupont, S. & Wickstrom, S.A. Mechanical regulation of chromatin and transcription. Nat Rev Genet 23, 624–643 (2022).

20. Chalut, K.J. et al. Label-free, high-throughput measurements of dynamic changes in cell nuclei using angle-resolved low coherence interferometry. Biophys J 94, 4948–4956 (2008).

21. Domingo-Muelas, A. et al. Human embryo live imaging reveals nuclear DNA shedding during blastocyst expansion and biopsy. Cell 186, 3166–3181 e3118 (2023).

22. Erickson, G.R., Alexopoulos, L.G. & Guilak, F. Hyper-osmotic stress induces volume change and calcium transients in chondrocytes by transmembrane, phospholipid, and G-protein pathways. J Biomech 34, 1527–1535 (2001).

23. Finan, J.D., Chalut, K.J., Wax, A. & Guilak, F. Nonlinear osmotic properties of the cell nucleus. Ann Biomed Eng 37, 477–491 (2009).

24. Skory, R.M. et al. The nuclear lamina couples mechanical forces to cell fate in the preimplantation embryo via actin organization. Nat Commun 14, 3101 (2023).

25. Keizer, V.I.P. et al. Live-cell micromanipulation of a genomic locus reveals interphase chromatin mechanics. Science 377, 489–495 (2022).

26. Maeshima, K., Iida, S. & Tamura, S. Physical Nature of Chromatin in the Nucleus. Cold Spring Harb Perspect Biol 13 (2021).

27. Nava, M.M. et al. Heterochromatin-Driven Nuclear Softening Protects the Genome against Mechanical Stress-Induced Damage. Cell 181, 800–817 e822 (2020).

28. Strickfaden, H. et al. Condensed Chromatin Behaves like a Solid on the Mesoscale In Vitro and in Living Cells. Cell 183, 1772–1784 e1713 (2020).

29. Muzzopappa, F., Hertzog, M. & Erdel, F. DNA length tunes the fluidity of DNA-based condensates. Biophys J 120, 1288–1300 (2021).

30. Schneider, M.W.G. et al. A mitotic chromatin phase transition prevents perforation by microtubules. Nature 609, 183–190 (2022).

31. Larson, A.G. & Narlikar, G.J. The Role of Phase Separation in Heterochromatin Formation, Function, and Regulation. Biochemistry 57, 2540–2548 (2018).

32. Rossant, J. & Tam, P.P.L. Early human embryonic development: Blastocyst formation to gastrulation. Dev Cell 57, 152–165 (2022).

33. Perera, M. & Brickman, J.M. In vitro models of human hypoblast and mouse primitive endoderm. Curr Opin Genet Dev 83, 102115 (2023).

34. De Belly, H. et al. Membrane Tension Gates ERK-Mediated Regulation of Pluripotent Cell Fate. Cell Stem Cell 28, 273–284 e276 (2021).

35. Chen, G. et al. Chemically defined conditions for human iPSC derivation and culture. Nat Methods 8, 424–429 (2011).

36. Chu, F.Y., Haley, S.C. & Zidovska, A. On the origin of shape fluctuations of the cell nucleus. Proc Natl Acad Sci U S A 114, 10338–10343 (2017).

37. Lomakin, A.J. et al. The nucleus acts as a ruler tailoring cell responses to spatial constraints. Science 370 (2020).

38. Clarke, D.N. & Martin, A.C. Actin-based force generation and cell adhesion in tissue morphogenesis. Curr Biol 31, R667–R680 (2021).

39. Szórádi, T. et al. nucGEMs probe the biophysical properties of the nucleoplasm. bioRxiv, 2021.2011.2018.469159 (2021).

40. Muncie, J.M. et al. Mechanical Tension Promotes Formation of Gastrulation-like Nodes and Patterns Mesoderm Specification in Human Embryonic Stem Cells. Dev Cell 55, 679–694 e611 (2020).

41. Valet, M., Siggia, E.D. & Brivanlou, A.H. Mechanical regulation of early vertebrate embryogenesis. Nat Rev Mol Cell Biol 23, 169–184 (2022).

42. Dattani, A., Huang, T., Liddle, C., Smith, A. & Guo, G. Suppression of YAP safeguards human naive pluripotency. Development 149 (2022).

43. Hashimoto, M. & Sasaki, H. Epiblast Formation by TEAD-YAP-Dependent Expression of Pluripotency Factors and Competitive Elimination of Unspecified Cells. Dev Cell 50, 139–154 e135 (2019).

44. Meyer, K., Lammers, N.C., Bugaj, L.J., Garcia, H.G. & Weiner, O.D. Optogenetic control of YAP reveals a dynamic communication code for stem cell fate and proliferation. Nat Commun 14, 6929 (2023).

45. Regin, M., et al. Lineage segregation in human pre-implantation embryos is specified by YAP1 and TEAD1. bioRxiv, 2022.2009.2029.509946 (2023).

46. Zhu, M. et al. Human embryo polarization requires PLC signaling to mediate trophectoderm specification. eLife 10, e65068 (2021).

47. Gerri, C. et al. Initiation of a conserved trophectoderm program in human, cow and mouse embryos. Nature 587, 443–447 (2020).

48. Petropoulos, S. et al. Single-Cell RNA-Seq Reveals Lineage and X Chromosome Dynamics in Human Preimplantation Embryos. Cell 165, 1012–1026 (2016).

49. Zhu, M. et al. Human embryo polarization requires PLC signaling to mediate trophectoderm specification. Elife 10 (2021).

50. Bravo Gonzalez-Blas, C., et al. SCENIC+: single-cell multiomic inference of enhancers and gene regulatory networks. Nat Methods 20, 1355–1367 (2023).

51. Schep, A.N., Wu, B., Buenrostro, J.D. & Greenleaf, W.J. chromVAR: inferring transcription-factor-associated accessibility from single-cell epigenomic data. Nat Methods 14, 975–978 (2017).

52. Schwalb, B. et al. TT-seq maps the human transient transcriptome. Science 352, 1225–1228 (2016).

53. Christoph, K., Beck, F.X. & Neuhofer, W. Osmoadaptation of Mammalian cells - an orchestrated network of protective genes. Curr Genomics 8, 209–218 (2007).

54. Murphy, L.O., MacKeigan, J.P. & Blenis, J. A network of immediate early gene products propagates subtle differences in mitogen-activated protein kinase signal amplitude and duration. Mol Cell Biol 24, 144–153 (2004).

55. Rivera, V.M. & Greenberg, M.E. Growth factor-induced gene expression: the ups and downs of c-fos regulation. New Biol 2, 751–758 (1990).

56. Coyle, P., Philcox, J.C., Carey, L.C. & Rofe, A.M. Metallothionein: the multipurpose protein. Cell Mol Life Sci 59, 627–647 (2002).

57. Hillmer, R.E. & Link, B.A. The Roles of Hippo Signaling Transducers Yap and Taz in Chromatin Remodeling. Cells 8 (2019).

58. Klopf, E., Schmidt, H.A., Clauder-Munster, S., Steinmetz, L.M. & Schuller, C. INO80 represses osmostress induced gene expression by resetting promoter proximal nucleosomes. Nucleic Acids Res 45, 3752–3766 (2017).

59. Hohfeld, J. et al. Maintaining proteostasis under mechanical stress. EMBO Rep 22, e52507 (2021).

60. Zhou, X., Naguro, I., Ichijo, H. & Watanabe, K. Mitogen-activated protein kinases as key players in osmotic stress signaling. Biochim Biophys Acta 1860, 2037–2052 (2016).

61. Watson, J.L. et al. Macromolecular condensation buffers intracellular water potential. Nature 623, 842–852 (2023).

62. Finan, J.D. & Guilak, F. The effects of osmotic stress on the structure and function of the cell nucleus. J Cell Biochem 109, 460–467 (2010).

63. Bracken, A.P., Dietrich, N., Pasini, D., Hansen, K.H. & Helin, K. Genome-wide mapping of Polycomb target genes unravels their roles in cell fate transitions. Genes Dev 20, 1123–1136 (2006).

64. Misteli, T. The Self-Organizing Genome: Principles of Genome Architecture and Function. Cell 183, 28–45 (2020).

65. Willemin, A., Szabo, D. & Pombo, A. Epigenetic regulatory layers in the 3D nucleus. Mol Cell 84, 415–428 (2024).

66. Eeftens, J.M., Kapoor, M., Michieletto, D. & Brangwynne, C.P. Polycomb condensates can promote epigenetic marks but are not required for sustained chromatin compaction. Nat Commun 12, 5888 (2021).

67. Plys, A.J. et al. Phase separation of Polycomb-repressive complex 1 is governed by a charged disordered region of CBX2. Genes Dev 33, 799–813 (2019).

68. Tatavosian, R. et al. Nuclear condensates of the Polycomb protein chromobox 2 (CBX2) assemble through phase separation. J Biol Chem 294, 1451–1463 (2019).

69. Jomova, K. et al. Essential metals in health and disease. Chem Biol Interact 367, 110173 (2022).

70. Sim, E.Z. et al. Methionine metabolism regulates pluripotent stem cell pluripotency and differentiation through zinc mobilization. Cell Rep 40, 111120 (2022).

71. Lu, J. et al. Single-cell RNA sequencing reveals metallothionein heterogeneity during hESC differentiation to definitive endoderm. Stem Cell Res 28, 48–55 (2018).

72. Slamecka, J. et al. Induced pluripotent stem cells derived from human amnion in chemically defined conditions. Cell Cycle 17, 330–347 (2018).

73. Lanner, F. & Rossant, J. The role of FGF/Erk signaling in pluripotent cells. Development 137, 3351–3360 (2010).

74. Mas, G. & Di Croce, L. The role of Polycomb in stem cell genome architecture. Curr Opin Cell Biol 43, 87–95 (2016).

75. Vining, K.H. & Mooney, D.J. Mechanical forces direct stem cell behaviour in development and regeneration. Nat Rev Mol Cell Biol 18, 728–742 (2017).

76. Schindelin, J., et al. Fiji: an open-source platform for biological-image analysis. Nat Methods 9, 676–682 (2012).

77. Machado, S., Mercier, V. & Chiaruttini, N. LimeSeg: a coarse-grained lipid membrane simulation for 3D image segmentation. BMC Bioinformatics 20, 2 (2019).

78. Pachitariu, M. & Stringer, C. Cellpose 2.0: how to train your own model. Nat Methods 19, 1634–1641 (2022).

79. Stringer, C., Wang, T., Michaelos, M. & Pachitariu, M. Cellpose: a generalist algorithm for cellular segmentation. Nat Methods 18, 100–106 (2021).

80. Ollion, J., Cochennec, J., Loll, F., Escude, C. & Boudier, T. TANGO: a generic tool for high-throughput 3D image analysis for studying nuclear organization. Bioinformatics 29, 1840–1841 (2013).

81. Legland, D., Arganda-Carreras, I. & Andrey, P. MorphoLibJ: integrated library and plugins for mathematical morphology with ImageJ. Bioinformatics 32, 3532–3534 (2016).

82. Le Berre, M., Aubertin, J. & Piel, M. Fine control of nuclear confinement identifies a threshold deformation leading to lamina rupture and induction of specific genes. Integr Biol (Camb) 4, 1406–1414 (2012).

83. Keikhosravi, A., et al. HiTIPS: High-Throughput Image Processing Software for the Study of Nuclear Architecture and Gene Expression. bioRxiv (2023).

84. Armiger, T.J., Lampi, M.C., Reinhart-King, C.A. & Dahl, K.N. Determining mechanical features of modulated epithelial monolayers using subnuclear particle tracking. J Cell Sci 131 (2018).

85. Keegan, S., Fenyö, D. & Holt, L.J. GEMspa: a Napari plugin for analysis of single particle tracking data. *bioRxiv*, 2023.2006.2026.546612 (2023).

86. Tirosh, I. et al. Single-cell RNA-seq supports a developmental hierarchy in human oligodendroglioma. Nature 539, 309–313 (2016).

87. Zhang, Y. et al. Model-based analysis of ChIP-Seq (MACS). Genome Biol 9, R137 (2008).

88. Tarasov, A., Vilella, A.J., Cuppen, E., Nijman, I.J. & Prins, P. Sambamba: fast processing of NGS alignment formats. Bioinformatics 31, 2032–2034 (2015).

89. Amemiya, H.M., Kundaje, A. & Boyle, A.P. The ENCODE Blacklist: Identification of Problematic Regions of the Genome. Sci Rep 9, 9354 (2019).

90. Quinlan, A.R. & Hall, I.M. BEDTools: a flexible suite of utilities for comparing genomic features. Bioinformatics 26, 841–842 (2010).

91. Fursova, N.A. et al. Synergy between Variant PRC1 Complexes Defines Polycomb-Mediated Gene Repression. Mol Cell 74, 1020–1036 e1028 (2019).

92. Ramirez, F. et al. deepTools2: a next generation web server for deep-sequencing data analysis. Nucleic Acids Res 44, W160–165 (2016).

93. Love, M.I., Huber, W. & Anders, S. Moderated estimation of fold change and dispersion for RNA-seq data with DESeq2. Genome Biol 15, 550 (2014).

94. Fursova, N.A. et al. BAP1 constrains pervasive H2AK119ub1 to control the transcriptional potential of the genome. Genes Dev 35, 749–770 (2021).

95. Chen, E.Y. et al. Enrichr: interactive and collaborative HTML5 gene list enrichment analysis tool. BMC Bioinformatics 14, 128 (2013).

96. Dobin, A. et al. STAR: ultrafast universal RNA-seq aligner. Bioinformatics 29, 15–21 (2013).

97. Tyanova, S., Temu, T. & Cox, J. The MaxQuant computational platform for mass spectrometry-based shotgun proteomics. Nat Protoc 11, 2301–2319 (2016).

98. Hornbeck, P.V. et al. PhosphoSitePlus, 2014: mutations, PTMs and recalibrations. Nucleic Acids Res 43, D512–520 (2015).

99. Nordin, A., Zambanini, G., Pagella, P. & Cantù, C. The CUT&RUN Blacklist of Problematic Regions of the Genome. bioRxiv, 2022.2011.2011.516118 (2022).

