## Supplementary Materials for "Mechano-osmotic signals control chromatin state and fate transitions in pluripotent stem cells"

This document contains Supplementary Figures (S1-S7)

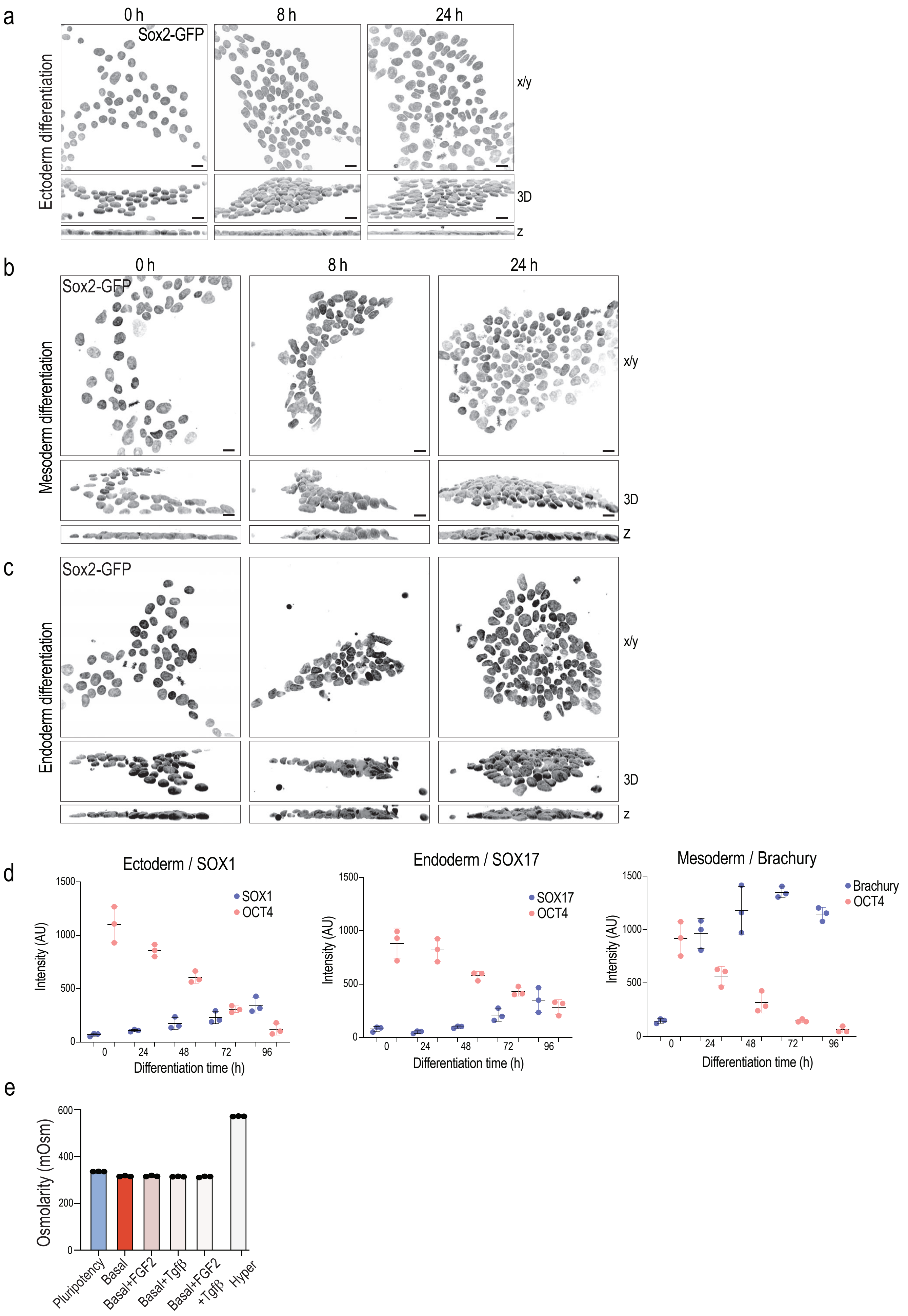


Supplementary Fig.1

**Supplementary Figure S1.**
**(a-c)** Representative top views (x-y), 3D reconstructions and cross sections (z), and quantification of nuclear height and volume from of Sox-GFP-tagged hiPSCs undergoing ectodermal (a), mesodermal (b), or endodermal (c) differentiation for the indicated time points. Note progressive decline in nuclear volume (scale bars 15 µm; n= 3 independent experiments with >330 cells/condition/experiment; ANOVA/ Dunnett's). **(d)** Quantification of lineage marker intensity from immunofluorescence stainings in hiPSCs undergoing ectodermal (left), endodermal (middle), or mesodermal (right) differentiation for the indicated time points (scale bars 15 µm; n= 3 independent experiments with >330 cells/condition/experiment). **(e)** Osmolarity measurements of the various media conditions (mean ± SD; n=3 independent measurements).

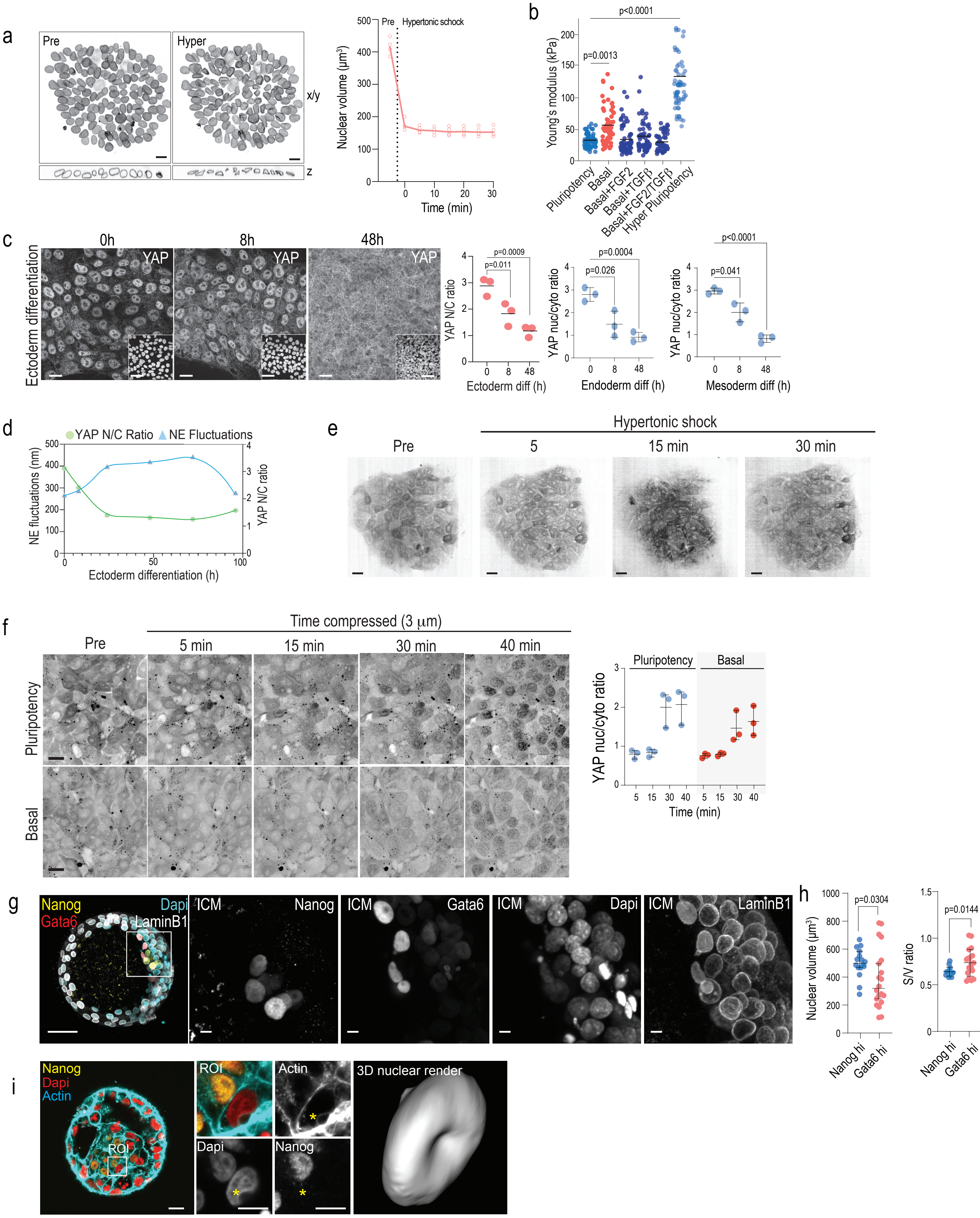


Supplementary Fig.2

**Supplementary Figure S2.**

**(a)** Representative snapshots of live imaging (x/y), optical cross sections (z), and quantifications from LaminB1-RFP-tagged hiPSCs before (Pre) and after (Hyper) hypertonic shock. Note progressive decline in nuclear volume. Line represents median volume and individual dots are average colony volumes at indicated timepoints (scale bars 10 µm; n=3 independent experiments with >80 cells/condition/experiment; Mann-Whitney). **(b)** AFM force indentation experiments of iPS cell nuclei within 20 min of media switch or hypertonic shock. Note data is reproduced from Fig. 1m but with additional condition of hypertonic shock (n = > 70 nuclei/condition pooled across 5 independent experiments; ANOVA/Kruskal-Wallis). **(c)** Quantification of YAP nuclear-to-cytoplasm (nuc/cyto) ratio from cells undergoing ectodermal, endodermal, or mesodermal differentiation at timepoints indicated. Note progressive decline in nuclear YAP (n= 3 independent experiments with >990 cells/experiment independent experiments; ANOVA/ Kruskal-Wallis). **(d)** Correlation plot of nuclear YAP and nuclear envelope fluctuations in iPSCs undergoing ectodermal differentiation for timepoints indicated. Note anticorrelation of nuclear YAP and fluctuations (representative of 3 independent experiments is shown). **(e)** Representative snapshots of live imaging of YAP-Halo-tag hiPSCs during hypertonic shock (scale bars 10 µm; n=3 independent experiments). **(f)** Representative snapshots of live imaging and quantification from YAP-Halo-tag hiPSCs during 3 µm compression in pluripotency or basal medium. Note comparable activation of YAP in both conditions (scale bars 30 µm; n=3 independent experiments with >280cells/condition/experiment). **(g)** Representative images and quantification of nuclear volume human preimplantation stage embryos stained for Gata6, Nanog, and LaminB1. **(h)** Nuclear volume of Gata6-high cells in the inner cell mass (scale bars 50 and 5 µm; n= 20 (Gata high) and 16 (Nanog high) nuclei pooled across 5 embryos; Mann Whitney). **(i)** Representative images of human preimplantation stage embryos stained for Dapi, Nanog and Actin (phalloidin). Note actin-rich bleb like structures with corresponding nuclear deformation (scale bars 20 and 10 µm; images representative of 3 embryos).

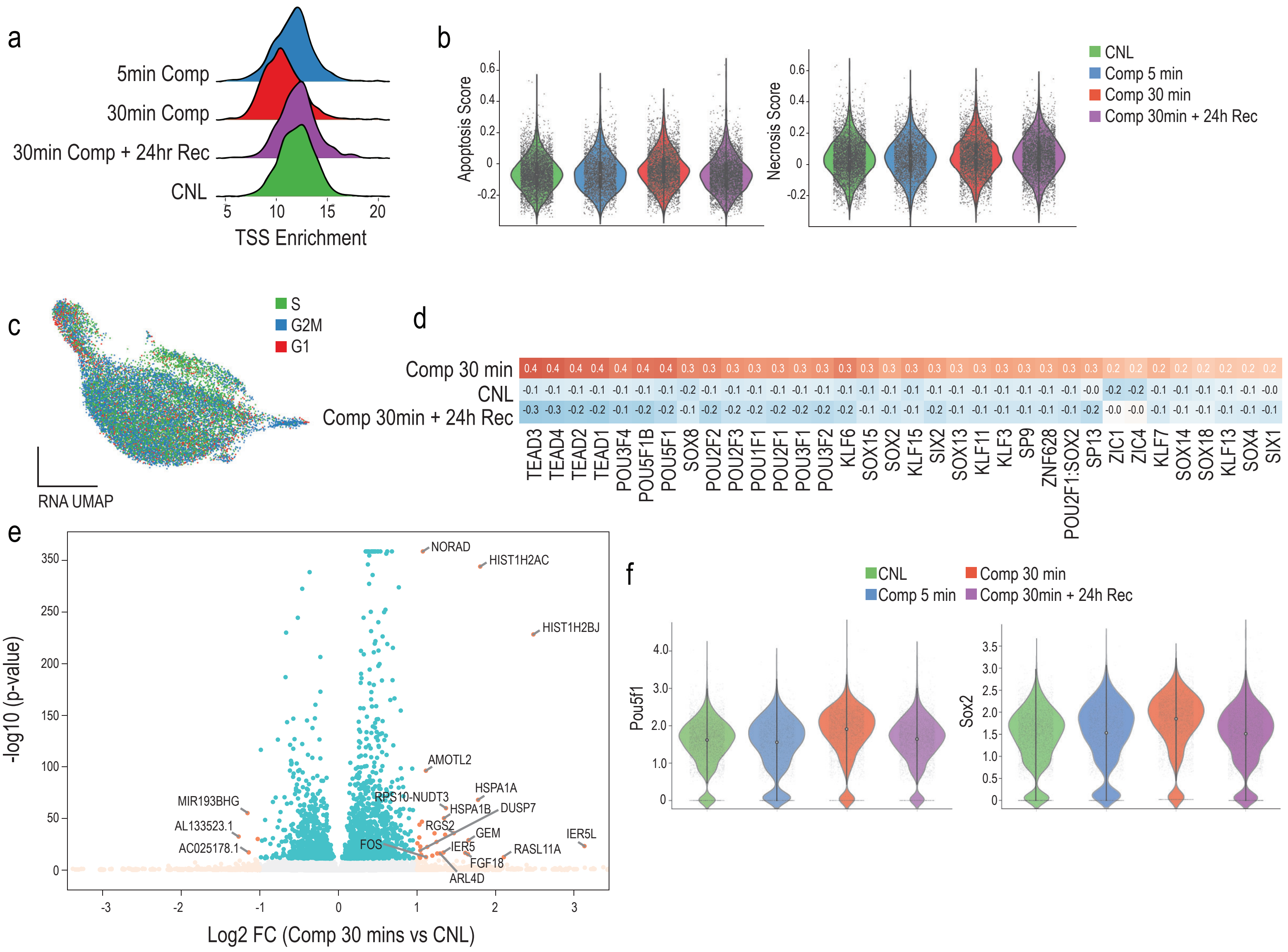


**Supplementary Figure S3.**
(a) Quantification of open chromatin from scATACseq of iPSCs compressed for 5 or 30 mins with or without 24 h recovery as well as uncompressed controls (CNL). Note reduced accessibility at transcription start sites (TSS) at 30 min compression. (b) Violin plots of computed apoptosis and necrosis scores computed from single cell RNAseq across all conditions. (c) UMAP of cell cycle stage distribution from single cell RNAseq across all conditions. (d) Heatmap of ChromVAR analysis from scATACseq of iPSCs compressed for 30 mins with or without 24 h recovery. (e) Volcano plot of differential gene expression between 30 min compression and CNL. (f) Violin plots of POU5F1 and SOX2 gene expression across all conditions.

a

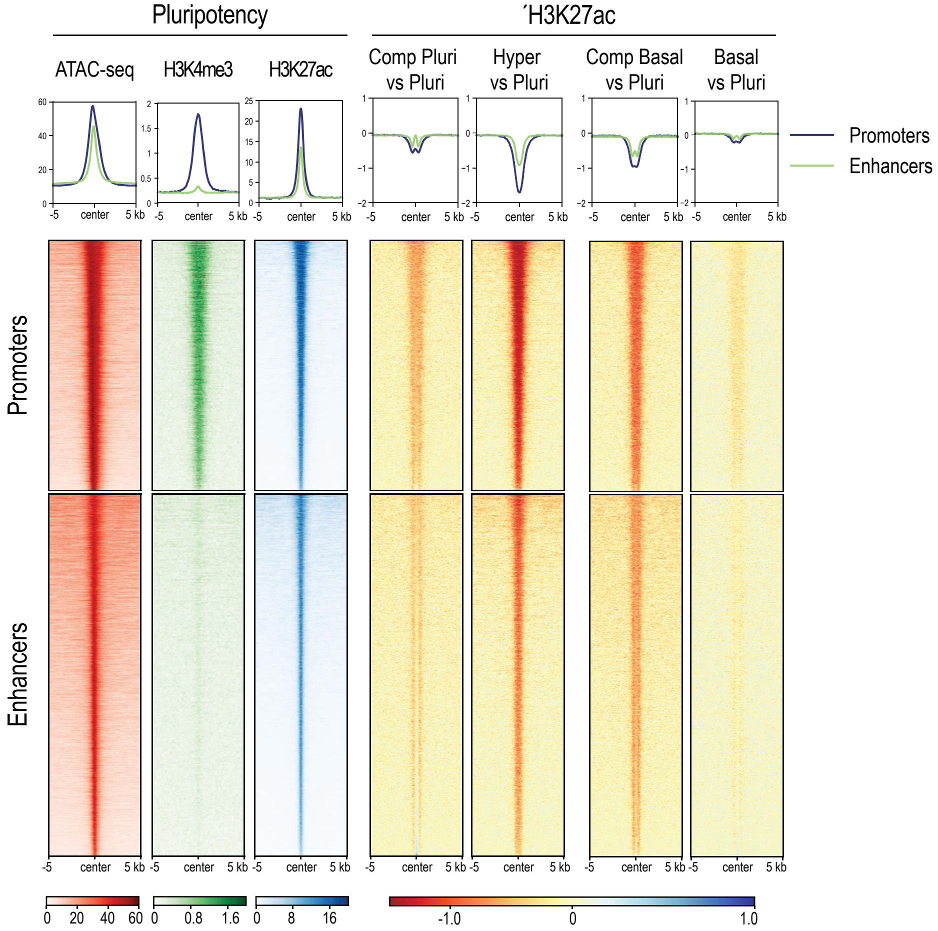


b

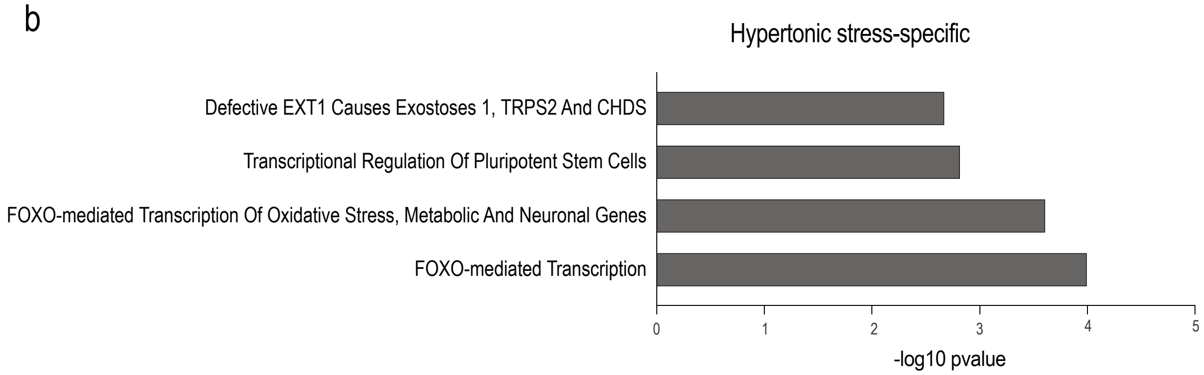


**Supplementary Figure S4.**
(a) Heatmaps of pseudo-bulk ATAC-seq, H3K4me3 and H3K27ac Cut & Tag at annotated H3K27ac peaks, further classified into active promoters and putative active enhancers, illustrating expected enrichment of histone modifications and chromatin accessibility at promoters and enhancers. Log2 fold change (FC) heatmaps highlight H3K27ac changes at active promoters and enhancers across different conditions compared to the pluripotency medium condition. (b) Reactome pathway enrichment of decommissioned enhancers specific to hypertonic shock.

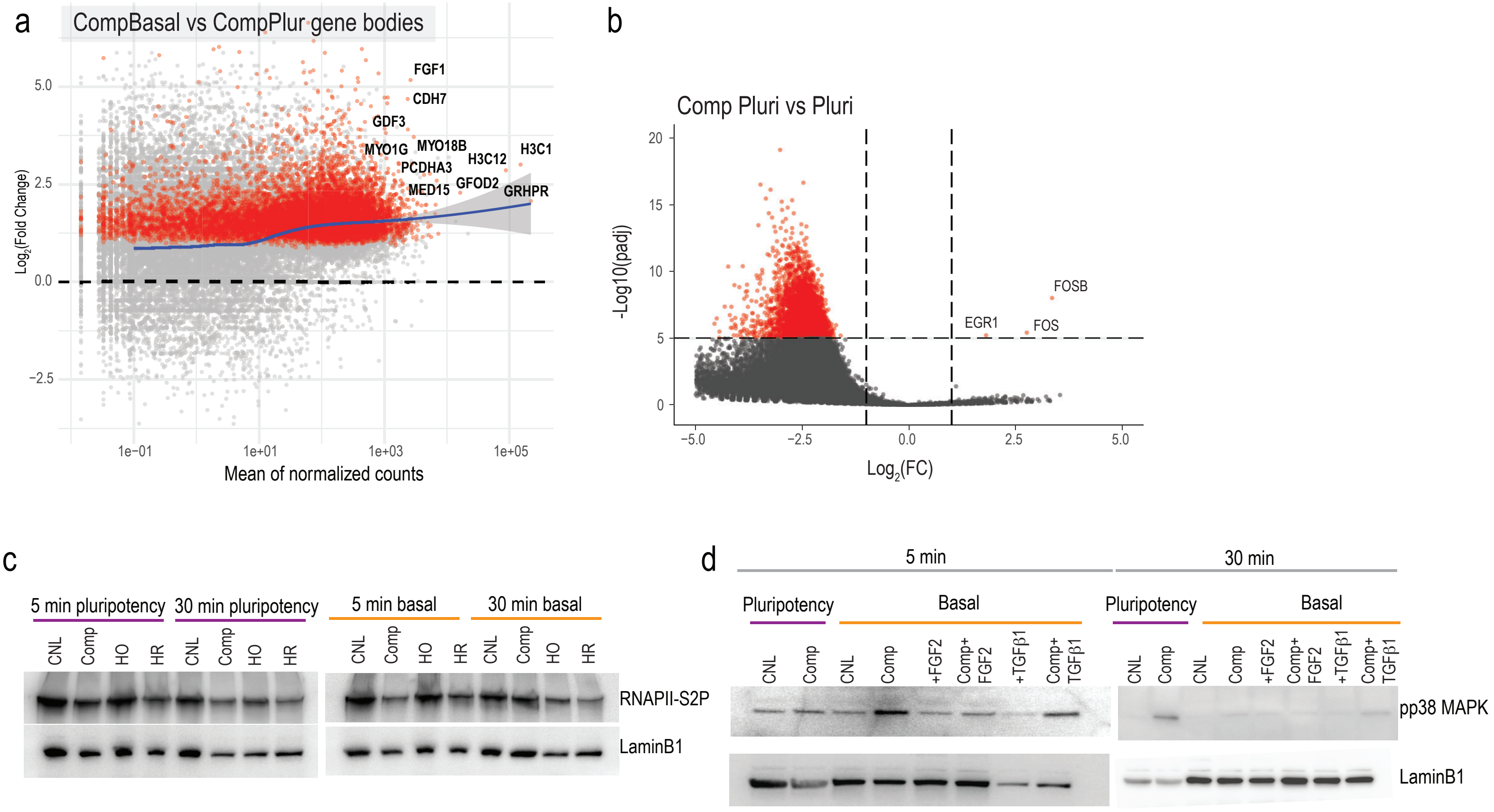


**Supplementary Figure S5.**

(a) Spike-in normalized MA plot contrasting CompBasal to CompPluri nascent transcript states. Loess local regression line in blue. Selected statistically significant upregulated genes are labeled. (b) Spike-in normalized volcano plot contrasting CompPluri to Pluri conditions. Statistically significant upregulated genes are labeled. (c) Representative western blots from RNAPII-S2P from cells exposed to compression or hypertonic/hypotonic shocks in the media compositions and time points indicated. Note decrease in RNAPII-S2P at 30 min compression in pluripotency medium but already at 5 min in basal medium (n=3 independent experiments). (d) Representative western blots from phosphorylated p38 (pp38) from cells exposed to compression in the media compositions and time points indicated. Note increase pf pp38 at 30 min compression in pluripotency medium but already at 5 min in basal medium and suppression of compression mediated activation of p38 by addition of FGF2 but not with TGF- into basal medium (n=3 independent experiments). Hypertonic (HR) and hypotonic (HO) shocks are used as controls for c, d.

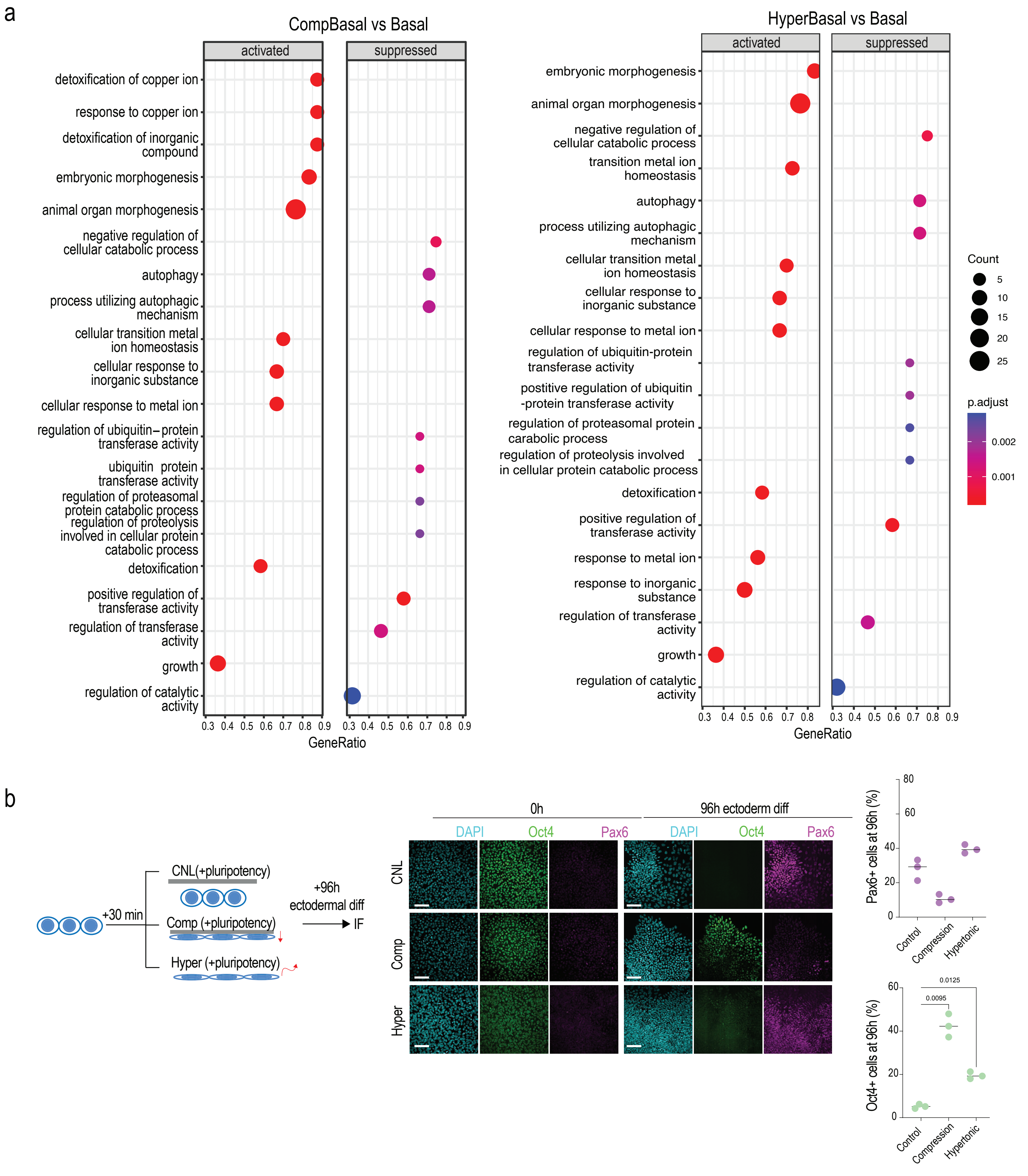


**Supplementary Figure S6.**
(a) Gene set enrichment analyses of differentially expressed genes from bulk RNA sequencing in 24 h basal medium versus 30 min compression + 24 h recovery in basal medium (left panel) and 24 h basal medium versus 30 min hypertonic shock + 24 h recovery in basal medium (right panel). Note enrichment of gene sets involved in morphogenesis and metal ion homeostasis in both conditions. (b) Schematic representation of experimental outline, representative images of Oct4 and Pax6 staining and quantification of cells exposed to 30 min compression or 30 min hypertonic shock in pluripotency medium followed by 96 h of ectodermal differentiation. Note delayed differentiation in both shock conditions (scale bars 75 µm; n=3 independent experiments with >800 cells/condition/experiment; ANOVA/Kruskal-Wallis).

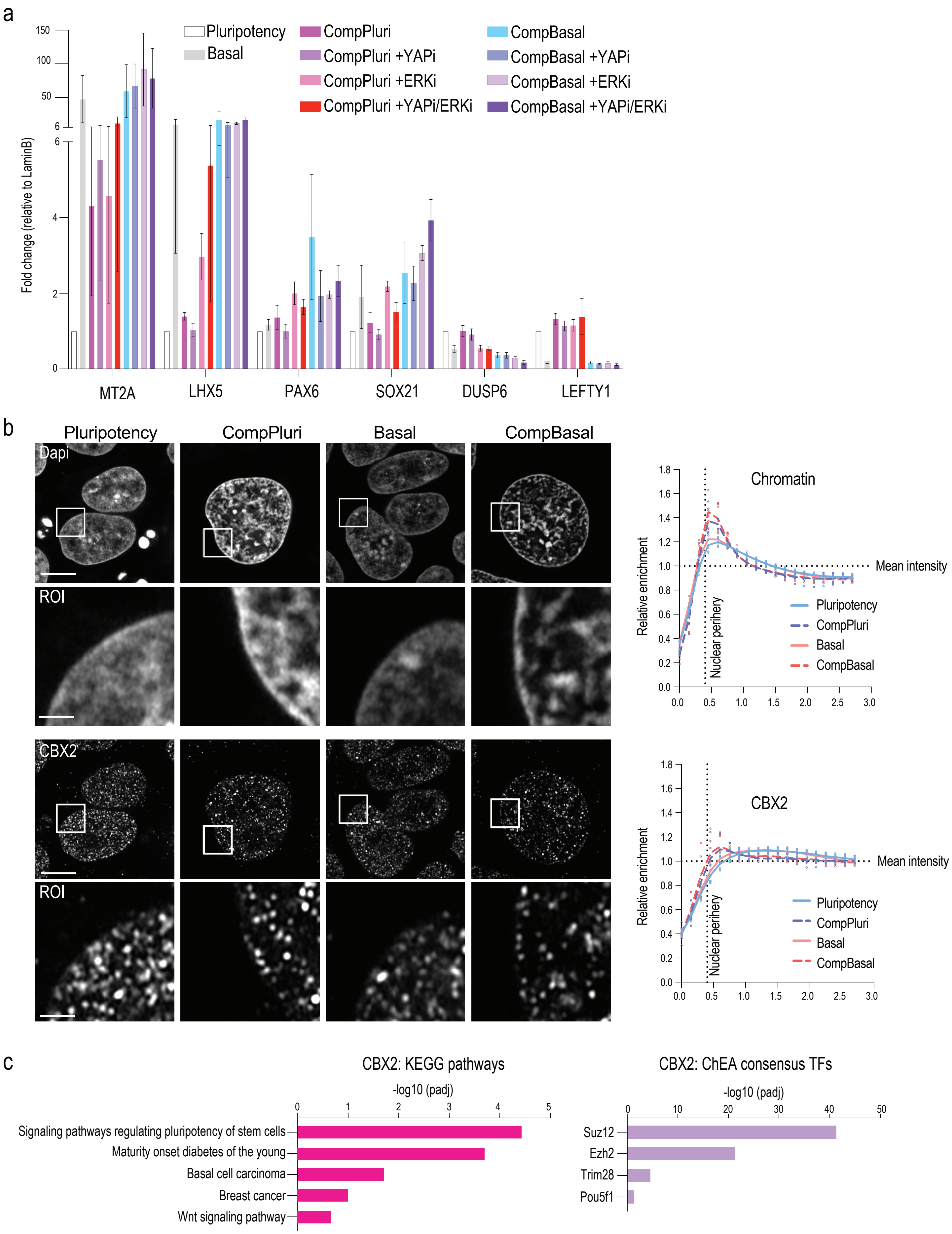
a
Fold change (relative to LaminB)
150
100
50
6
6
4
2
0
Pluripotency
Basal
CompPluri
CompPluri +YAPi
CompPluri +ERKi
CompPluri +YAPi/ERKi
CompBasal
CompBasal +YAPi
CompBasal +ERKi
CompBasal +YAPi/ERKi
MT2A
LHX5
PAX6
SOX21
DUSP6
LEFTY1
b
Pluripotency
CompPluri
Basal
CompBasal
Dapi
ROI
CBX2
ROI
Chromatin
Relative enrichment
Mean intensity
Nuclear perihery
Pluripotency
CompPluri
Basal
CompBasal
1.8
1.6
1.4
1.2
1.0
0.8
0.6
0.4
0.2
0.0
0.0
0.5
1.0
1.5
2.0
2.5
3.0
CBX2
Relative enrichment
Mean intensity
Nuclear perihery
Pluripotency
CompPluri
Basal
CompBasal
c
CBX2: KEGG pathways
-log10 (padj)
0
1
2
3
4
5
Signaling pathways regulating pluripotency of stem cells
Maturity onset diabetes of the young
Basal cell carcinoma
Breast cancer
Wnt signaling pathway
CBX2: ChEA consensus TFs
-log10 (padj)
0
10
20
30
40
50
Suz12
Ezh2
Trim28
Pou5f1

**Supplementary Figure S7.**

**(a)** RT-qPCR from cells exposed to 30 min compression in indicated media conditions followed by 24h recovery in the same media conditions with and without YAP and ERK inhibitors as indicated. Note increased metallothionine (MT2A) and differentiation gene (LHX5, PAX6, SOX21) expression in basal medium and 30 min compression in basal medium conditions (CompBasal) and lack of rescue with YAP/ERK inhibition. Cells compressed in pluripotency medium (CompPluri) show increased differentiation gene expression upon inhibition of YAP/ERK (mean ± SD; n=3 independent experiments). **(b)** Representative super resolution images and quantification of CBX2 clustering at the nuclear periphery. Scale bars 5 µm and 10 µm for ROIs (n=5 independent experiments with >20 cells/condition/experiment). **(c)** KEGG pathway analysis (left) and CheEA consensus Transcription Factor (TF) prediction (right) from CBX2 peaks differentially occupied in basal/compression in basal medium conditions.
